# Integrative analysis of the 3D genome and epigenome in mouse embryonic tissues

**DOI:** 10.1101/2022.04.25.489471

**Authors:** Miao Yu, Nathan R. Zemke, Ziyin Chen, Ivan Juric, Rong Hu, Ramya Raviram, Armen Abnousi, Rongxin Fang, Yanxiao Zhang, David U. Gorkin, Yang Li, Yuan Zhao, Lindsay Lee, Anthony D. Schmitt, Yunjiang Qiu, Diane E. Dickel, Axel Visel, Len A. Pennacchio, Ming Hu, Bing Ren

## Abstract

While a rich set of putative *cis*-regulatory sequences involved in mouse fetal development has been annotated recently based on chromatin accessibility and histone modification patterns, delineating their role in developmentally regulated gene expression continues to be challenging. To fill this gap, we mapped chromatin contacts between gene promoters and distal sequences genome-wide in seven mouse fetal tissues, and for one of them, across six developmental stages. We identified 248,620 long-range chromatin interactions centered at 14,138 protein-coding genes and characterized their tissue-to-tissue variations as well as developmental dynamics. Integrative analysis of the interactome with previous epigenome and transcriptome datasets from the same tissues revealed a strong correlation between the chromatin contacts and chromatin state at distal enhancers, as well as gene expression patterns at predicted target genes. We predicted target genes of 15,098 candidate enhancers, and used them to annotate target genes of homologous candidate enhancers in the human genome that harbor risk variants of human diseases. We present evidence that schizophrenia and other adult disease risk variants are frequently found in fetal enhancers, providing support for the hypothesis of fetal origins of adult diseases.

## Introduction

Developmental programs in multicellular organisms involve temporal and spatially controlled expression of genes, which are driven by the dynamic binding of sequence-specific transcription factors to *cis*-regulatory elements^1–4^. Recently, the Encyclopedia of DNA Elements (ENCODE)^5, 6^ project annotated a rich set of candidate *cis*-regulatory elements (cCREs) in the mouse genome through comprehensive mapping of histone modifications, chromatin accessibility and DNA methylation across a wide spectrum of tissues and early developmental stages^7, 8^. While these catalogs of cCREs have revealed the location and tissue-specific usage of regulatory sequences in the genome, understanding their role in developmental programs remains a daunting challenge, owing to our incomplete knowledge of the regulatory target genes of most cCREs.

The traditional way of assigning the target genes of cCREs to its closest promoter in one-dimensional (1D) genomic distance does not take into account the fact that cCREs frequently regulate genes over a large 1D genomic distance by skipping over closer genes^9^. Indeed, active cCREs (e.g. enhancers) and their target gene promoters are believed to be spatially close in three-dimensional (3D) space. Consequently, high-throughput chromatin conformation capture based techniques, such as Hi-C^10, 11^, capture Hi-C^12, 13^, chromatin interaction analysis by paired-end tag (ChIA-PET)^14, 15^ and proximity ligation-assisted ChIP-seq (PLAC-seq and HiChIP)^16, 17^, have been increasingly used to map long-range chromatin interactions between promoters and cCREs to infer target genes of cCREs. Multiple large-scale studies have characterized long-range chromatin interactions across a wide spectrum of human and mouse tissues (or cell types)^18–23^. It has been demonstrated that tissue-specific interactions are not only correlated with changes in gene expression, but also enriched for tissue-specific cCREs and noncoding risk variants associated with diseases, revealing the power of using 3D genome information to identify the function of cCREs and their contribution to disease etiology. However, the majority of previous studies focused on adult tissues, and the chromatin interactomes in developing fetus have yet to be systematically investigated.

In this work, we mapped chromatin interactions centered at 14,138 protein-coding gene promoters in mouse embryos, spanning seven tissues and, for one of them, across six developmental stages. To investigate the role of CTCF in mediating the promoter-centered interactions, we also generated CTCF ChIP-seq datasets using the same samples. Integrating with previous epigenomic profiles from the same tissues and developmental stages, representing 8 histone modifications (H3K4me3, H3K4me2, H3K4me1, H3K27ac, H3K9ac, H3K36me3, H3K27me3 and H3K9me3), chromatin accessibitliy^7^ and gene expression^24^ landscapes^5^, we examined the relationships between long-range chromatin interactions, chromatin states, chromatin accessibility, CTCF binding and gene expression during mouse fetal development. We predicted enhancer target genes and explained tissue-specific and developmental stage restricted gene expression programs. We further demonstrated that 3D genome information may facilitate the functional annotation of noncoding risk variants in the human genome and prioritization of gene targets for complex human diseases.

### Mapping promoter-centered chromatin interactome in mouse fetal tissues

We performed PLAC-seq experiments (also known as HiChIP) using antibodies against histone 3 lysine 4 trimethylation (H3K4me3), a chromatin mark found at both active and poised promoter regions, to map chromatin interactions centered at gene promoters. Seven mouse fetal tissues from embryonic day 12.5 (E12.5) were assayed in duplicates, including forebrain (FB), midbrain (MB), hindbrain (HB), neural tube (NT), limb (LM), craniofacial prominence (CF) and liver (LV). Forebrain tissues at five extra time points from E13.5 until birth (E13.5, E14.5, E15.5, E16.5 and postnatal day 0, P0) were further investigated (**Fig. 1A**). Two biological replicates were collected and assayed for each tissue, with a median sequencing depth of 206 million reads per replicate. Each replicate was subjected to a set of quality control criteria to ensure the consistent data quality across different samples: 1) >60% intra-chromosomal (*cis*) reads; 2) >50% intra-chromosomal (*cis*) reads over 1 Kb distance and 3) >7.5% Fraction of Reads in Peaks (FRiP)^25^ (**Table S1**, see **Methods** for details).

**Figure 1.**
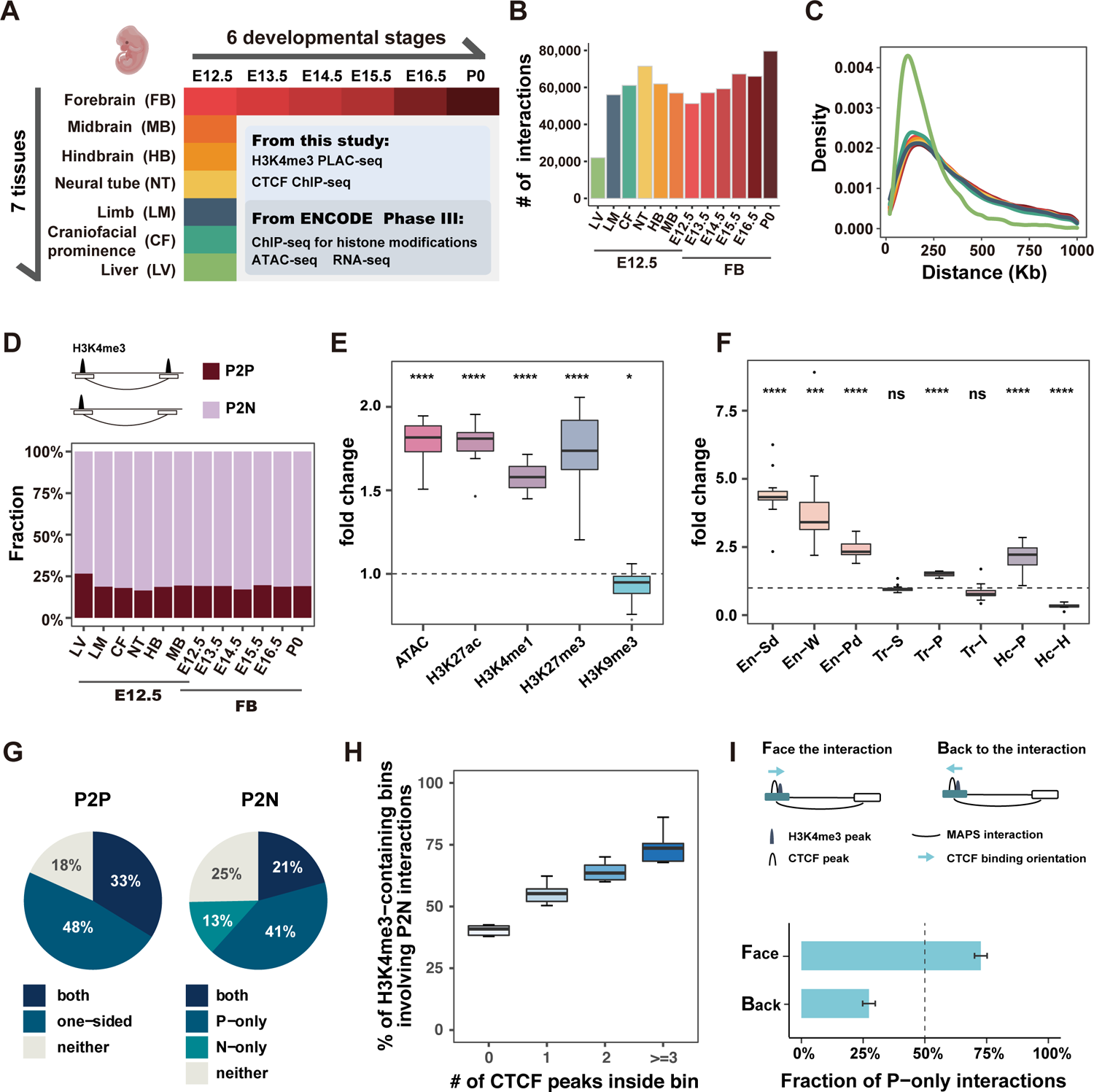
Characteristics of promoter-centered interactions identified from H3K4me3 PLAC-seq across 12 tissues and developmental stages during mouse fetal development. **A.** Overview of the experimental design. **B.** Number of MAPS-identified chromatin interactions for each tissue from the combined, down-sampled dataset. **C.** The density plot showed distribution of interaction distance for 12 tissues. Different tissues are represented by the colored schemes shown in Fig.1B. **D.** Fraction of Promoter-to-Promoter (P2P, red) and Promoter-to-Non-Promoter (P2N, pink) interactions across 12 samples. P2P interactions are defined as interactions with both anchor bins overlapping the reproducible H3K4me3 peaks identified in the same tissue; P2N interactions are defined as those with only one anchor bin overlapping the reproducible H3K4me3 peaks identified in the same tissue (as shown in the schematic diagram on the top). **E.** Boxplots showing the enrichment of promoter-interacting regions for accessible chromatins and histone marks of P2N interactions. Bins with equidistant from but on the other side of H3K4me3-containing bins were used as the control set. Fold-change of 1 is marked by the horizontal dashed line (n=12). *p <= 0.05, ****p <= 0.0001, from single sample student t-test comparing to mu= 1. **F.** Boxplots showing the enrichment of promoter-interacting regions within different chromatin states of P2N interactions. Bins with equidistant from but on the other side of H3K4me3-containing bins were used as the control set. En-Sd, Strong TSS-distal enhancer; En-W, Weak TSS-distal enhancer; En-Pd, Poised TSS-distal enhancer; Tr-S, Strong transcription; Tr-P, Permissive transcription; Tr-I, Initiation transcription; Hc-P, Polycomb-associated heterochromatin; Hc-H, H3K9me3-associated heterochromatin. Fold-change of 1 is marked by the horizontal dashed line. (n=12). ‘ns’: not significant, *p <= 0.05, ***p <= 0.001, ****p <= 0.0001, from single sample student t-test comparing to mu= 1. **G.** Average proportion of long-range interactions classified according to presence or absence of CTCF binding on one or both ends across 12 tissues (n=12). Promoter-to-Promoter (P2P) and Promoter-to-Non-Promoter (P2N) interactions are considered separately. P-only: CTCF binding on the “Promoter” end (10 Kb bins with H3K4me3 peak) only; N-only: CTCF binding on the “Non-Promoter” end (10 Kb bins without H3K4me3 peak) only. **H.** Boxplots showing the fraction of H3K4me3-containing 10 Kb bins with different number of CTCF peaks in the bin forming P2N interactions (n=12). **I.** A bar plot showing the fraction of P2N interactions formed on CTCF-anchored promoters (bins containing both H3K4me3 and CTCF ChIP-seq peaks) on each direction. Fraction of 0.5 is marked by the vertical dashed line, which shows the expected value from the random shuffle test (See Methods for details). Face: The orientation of CTCF binding motif on the Promoter end faces the direction of the Non-Promoter end; Back: The orientation of CTCF binding motif on the promoter end is back to the direction of the Non-Promoter end, as shown in the schematics on the top. The heights of the bars represent the average values of 12 tissues and the error-bars represent the standard deviation. For all boxplots in this figure: Horizontal line, median; box, interquartile range (IQR); whiskers, the most extreme value within ±1.5 × IQR.

To check the reproducibility between biological replicates, we first used the model-based analysis of PLAC-seq and HiChIP (MAPS) pipeline^26^ to identify statistically significant long-range chromatin interactions from each biological replicate at 10-Kb resolution. We found 54%-71% of interactions were shared between two biological replicates (**Fig. S1A**), which is at a similar level as in our previous study using mouse embryonic stem cells (mESCs)^26^. Additionally, the normalized contact frequencies at the reproducible interactions were also highly correlated with Pearson correlation coefficients ranging 0.77-0.80 (**Fig. S1B**). We therefore combined the data from two replicates of each tissue to increase the sensitivity of contacts mapping. To examine the dynamics in chromatin interactions across different tissues, we further down-sampled the combined data so that all 12 tissue/time point combinations had the same number of usable reads (see **Methods** for details). After re-applying MAPS to these sample-balanced datasets, a total of 248,620 interactions were identified at 10-Kb resolution across all tissues between 20 Kb to 1 Mb genomic distance. On average, ∼60,000 (51,264∼79,677, median 61,119) interactions were identified in each tissue excluding liver E12.5, where only 21,945 interactions were found (**Fig. 1B**). The median genomic distances that these interactions span were 270-310 Kb for eleven of the tissues, whereas in liver E12.5 the median distance was significantly shorter (170 Kb) (**Fig. 1C**). Approximately 20% (16%-27%, median 19%) of the identified interactions were between two H3K4me3-marked regions (hereinafter referred as “promoter-to-promoter” interactions, or P2P interactions), whereas the remaining interactions (73%-84%, median 81%) were between a genomic region associated with H3K4me3 and another one displaying no H3K4me3 signals (hereinafter referred as “Promoter-to-Non-promoter” interactions or P2N interactions) (**Fig. 1D**).

To evaluate the sensitivity of H3K4me3 PLAC-seq in detecting promoter-centered interactions, we compared our MAPS-identified interaction lists with published ones centered on 446 gene loci during mouse limb development revealed by Capture-C^27^. 77%∼85% (median 81%) of the top 1% Capture-C interactions (above the 99^th^ percentile) were recapitulated by H3K4me3 PLAC-seq experiments from the most closely related tissues, in comparison with only 4%∼11% (median 9%) recovery rate by distance-matched control sets (**Fig. S1C**). Even after lowering the threshold to the 95^th^ percentile for Capture-C interactions, more than 50% of them can still be found from PLAC-seq data (**Fig. S1D**).

### P2N interactions frequently occur between putative enhancers and promoters

As demonstrated in **Fig. 1D**, the majority of the identified interactions (>80%) were P2N interactions, indicating these interactions may link promoters (i.e. the H3K4me3-marked regions from PLAC-seq data) to their potential regulatory elements. Indeed, across all 12 tissues on average 38% of those promoter-interacting regions in P2N interactions contained accessible chromatin regions, defined by ATAC-seq peaks^7^, and such proportion (i.e., 38%) was significantly (P-value = 1.89e-10, one sample T-test comparing fold-change to mu=1) higher than the distance-matched control regions (21%) (**Fig. 1E, Fig. S2A**). Further exploration of their chromatin properties using ChIP-seq data of histone modifications^28^ suggested that they were likely enhancers: both enhancer markers, namely H3K4me1 (47% in P2N vs. 30% in control, P-value = 5.62e-11) and H3K27ac (26% in P2N vs. 15% in control, P-value = 2.41e-10), were enriched at these regions. The occurrence of H3K27me3 marks at these regions was also significantly higher than that in control regions (17% vs. 10%, P-value = 4.42e-07), consistent with previous reports of Polycomb-associated chromatin interactions^28–30^. In contrast, the heterochromatin mark H3K9me3 was depleted (6% vs. 7%, P-value = 0.015) (**Fig. 1E, Fig. S2A**). In line with above observations, integrative analysis with the 15-state ChromHMM model^31^ revealed 2-9 fold enrichment of all three types of distal enhancers (strong, weak and poised) and depletion of H3K9me3-associated heterochromatin at these non-promoter regions (**Fig. 1F, Fig. S2B**). Taken together, our data suggest that during mouse fetal development, promoter-centered chromatin interactions frequently occur between promoters and putative enhancers.

### CTCF is broadly involved in promoter-centered chromatin interactions during fetal development

CTCF plays a critical role in the formation of topological associating domains (TADs) through a cohesin-driven loop extrusion process^32^, but it is still not clear to what extent it is involved in chromatin organization at promoters during embryonic development. To investigate how much CTCF binding contributes to the organization of promotor-centered interactome in mouse fetal tissues, we performed chromatin immunoprecipitation with sequencing (ChIP–seq) assays for CTCF in all 12 tissue/time point combinations (**Fig. S3**). We found on average 82% of P2P interactions (75% by chance, P-value = 3.1e-7, paired T-test, see **Methods** for details) and 75% of P2N interactions (55% by chance, P-value = 1.9e-11, paired T-test) have CTCF binding on at least one end, supporting a general role for CTCF in the formation of promoter-centered interactions (**Fig. 1G**). In addition, 33% of P2P interactions and 21% of P2N interactions had CTCF binding on both ends (**Fig. S4A-B**), and approximately 50% of these “CTCF-CTCF” interactions exhibited convergent motif orientation, consistent with the loop extrusion model^33, 34^ (**Fig. S4C-D**).

Interestingly, a large proportion (54%) of P2N interactions have CTCF binding on only one end. Further classification based on the colocalization of CTCF with H3K4me3 showed that CTCF was more likely to be present on the promoter side of the interaction rather than the putative enhancer side (41% vs. 13%), suggesting a potential role for promoter-proximal CTCF binding in promoting the formation of promoter-enhancer interactions, a phenomenon that we previously reported for mESCs^35^. Consistent with this hypothesis, promoters associated with more CTCF binding sites nearby were also more likely to be involved in P2N interactions (**Fig. 1H, Fig. S4E-G**).

An earlier study from B cells posited that CTCF-binding regions can “reel in” other elements over long 1D genomic distances through cohesin-mediated loop extrusion, and therefore facilitate the formation of long-range interactions^36^. We hypothesized that the same mechanism also applies in mouse fetal tissues and promoter-proximal CTCF helps “reel in” genomic DNA in a specific orientation to search for distal putative enhancers for contact. If so, we expected that the orientation of promoter-proximal CTCF binding motifs would preferentially face the direction of promoter-interacting regions in P2N interactions. Indeed, among all P2N interactions with CTCF binding only on the promoter side, ∼75% had a CTCF binding motif facing its promoter-interacting region, compared to the random 50% chance (P-value = 1.4e-11, paired T-test) (**Fig. 1I**, see **Methods** for details). Collectively, these results support the role of CTCF in facilitating promoter-centered interactions in different tissues during mouse embryonic development.

### Promoter-centered interactome shows tissue-to-tissue variability and developmental dynamics that correlate with gene expression

To characterize the spatial and temporal dynamics of the promoter-centered interactomes during mouse fetal development, we performed principal component analysis using normalized PLAC-seq contact frequencies (see details in **Methods**) and found that tissues with similar lineages or developmental stages are clustered more closely (**Fig 2A**), consistent with the hierarchical clustering results based on H3K4me3 PLAC-seq data and H3K27ac ChIP-seq data (**Fig S5A-B**). We then evaluated the variability of promoter-centered interactomes across 7 tissues at E12.5, and asked how the tissue-specific chromatin interactions are associated with tissue-specific epigenome and gene expression. For this purpose, we selected a total of 15,027 P2N interactions that were identified only in the neural tissues (including FB, MB, HB and NT), in limb, in craniofacial or in liver, respectively (10.1% of all P2N interactions for tissues at E12.5, see details in **Methods**). We found that the normalized contact frequency of these neural-, limb-, craniofacial- and liver-restricted P2N interactions correlated strongly with gene expression levels and H3K27ac signals on the promoter-distal elements (**Fig 2B**). Moreover, we compared the promoter-centered interactomes between each pair of tissues at E12.5 and found that the difference in the number of interactions for each promoter is positively correlated with the changes of gene expression of the corresponding tissues (**Fig 2C, Fig S5C**). This trend is particularly pronounced between tissues that are of different developmental origins. For example, several neural-specific interactions were found at the promoter of *Myt1l,* a gene that plays a key role in neuronal differentiation^37^. Regions interacting with the *Myt1l* promoter showed higher H3K27ac in the forebrain at E12.5 compared to the other three tissues. We also observed limb-specific, craniofacial-specific and liver-specific interactions centered at the promoter of *Ppp1r3b* (a regulator of glycogen synthesis^38^), *Bves* (a gene encoding blood vessel/epicardial substance which may play a role in skeletal muscle development^39^) and *Rbp4* (a retinol carrier that delivers retinol from liver to peripheral tissues^40^) respectively, with high levels of H3K27ac on the promoter-interacting regions in the corresponding tissues (**Fig 2D**).

**Figure 2.**
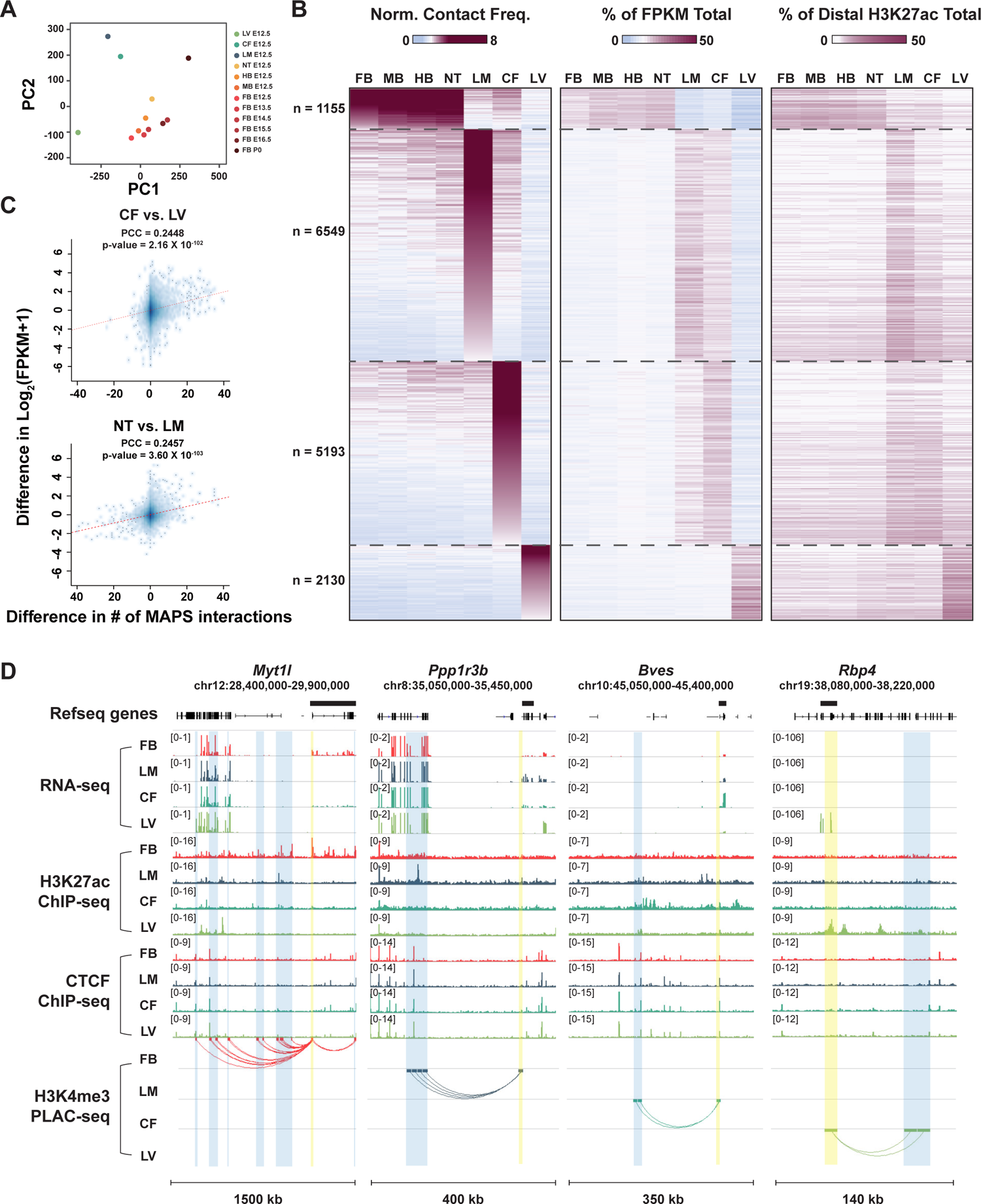
Tissue-to-tissue variability and developmental dynamics of promoter-centered interactomes. **A.** Principal component analysis (PCA) for the normalized PLAC-seq contact frequency. The first principal component (i.e., PC1, x-axis) accounts for 18.68% of the total variation, capturing the difference between 9 brain-related tissues and 3 non-brain-related tissues (LV, LM and CF). The second principal component (i.e., PC2, y-axis) accounts for 10.72% of the total variation, capture the difference between dynamic developmental stages. **B.** Heatmap displaying normalized contact frequencies, gene expression of interacting promoters, and H3K27ac distal peak signal in peak to non-peak tissue-specific interactions. Gene expression and H3K27ac are represented as individual tissue percentage of total sum across all tissues. Tissue-specific interactions are clustered for being exclusive in all brain- and NT, LM, CF, or LV. **C.** Scatterplot between the difference of the number of MAPS-identified significant chromatin interactions (x-axis) and the different of gene expression (y-axis, merged by log2(FPKM+1)). The top panel and the bottom panel display the comparison between CF and LV, and between NT and LM, respectively. The red dashed line represents the fitted linear line, suggesting that the change of significant chromatin interactions is positively correlated with the change of gene expression. **D.** IGV visualization of MAPS-identified interactions anchored at TSS regions around *Myt1l*, *Ppp1r3b, Bves* and *Rbp4* genes in tissues at embryonic day 12.5. Black boxes above the refseq gene track mark the gene boundary of each anchored genes. The bins containing H3K4me3 peaks (around TSS) are highlighted by yellow boxes (chr12: 29,520,000-295,400,00 for *Myt1l*; chr8: 35,370,000-35,380,000 for *Ppp1r3b*; chr10: 45,330,000-45,340,000 for *Bves*; chr19: 38,120,000-38,130,000 for *Rbp4*). MAPS-identified interactions overlapping these H3K4me3-containing bins are marked by arcs and the other ends of the interactions are highlighted by blue boxes. FB, forebrain. LM, limb. CF, craniofacial prominence. LV, liver.

We also examined the dynamics across developmental stages in the forebrain. We selected the E12.5/E13.5 FB-, E14.5/E15.5 FB-, and E16.5/P0 FB-restricted P2N interactions and performed a similar analysis as for tissue-restricted P2N interactions at E12.5 (**Fig S6A-B**). A positive correlation between interaction strength and gene expression was also observed although weaker than that between different tissues at E12.5, suggesting that the developmental dynamics of promoter interactome in general is less pronounced than the tissue-to-tissue variations. Taken together, our data demonstrate that the tissue-to-tissue variability and developmental dynamics of promoter-centered interactome are correlated with the fetal gene expression programs.

### Dynamic chromatin contacts at promoters coupled with chromatin state maps predict target genes of promoter-distal candidate enhancers

Long-range interactions between promoters and distal enhancers are believed to contribute to gene expression. However, a recent comprehensive study covering 24 diverse human cell types^23^ found that the variations in long-range interactions were only modestly correlated with the variations in gene expression, suggesting a complex relationship between the physical contacts between cCRE-gene pairs and gene expression. Recently, success has been achieved by combining the 3D chromatin information and chromatin state to predict the functional contributions of enhancers in gene expression^41, 42^. To assign candidate enhancers to their target genes in mouse fetal tissues, we leveraged the P2N interactions identified from H3K4me3 PLAC-seq data. Open chromatin regions in the interacting non-promoter bins defined by ATAC-seq peaks were assigned to the active or poised promoters (**Fig. 3A**). Only using the chromatin interactions, open chromatin regions, and H3K4me3 ChIP-seq peaks from the same sample, this integrative analysis identified 91,451 open chromatin sites to be in long-range chromatin with active or poised promoters of protein-coding genes, designated as interacting cCRE-gene pairs (**Fig. S7A**).

**Figure 3.**
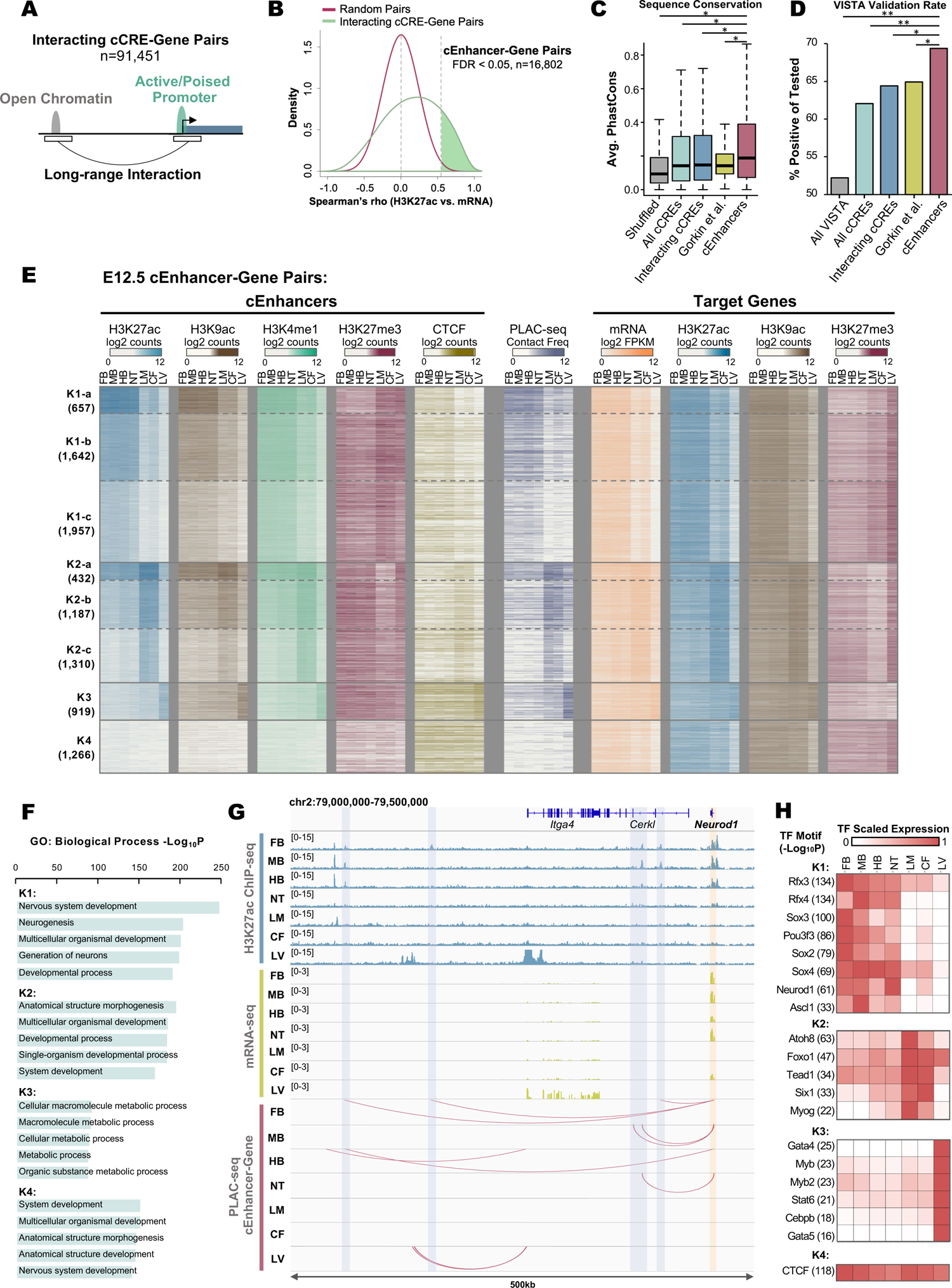
Profiling candidate enhancer-gene pairs in different fetal tissues. **A.** Schematic for assigning cCREs to target genes. **B.** Density plot of Spearman’s correlation coefficients between cCRE H3K27ac and interacting gene mRNA. **C.** Box plots of distributions for average Phast Cons scores for each nucleotide of all cCREs (d-TACs (Gorkin et al. 2020)^7^ with a chromatin accessible peak present in any of the 12 samples where PLAC-seq data was generated), size-matched random regions (shuffled), interacting cCREs (cCREs involved in 91,451 cCRE-gene pairs), predicted enhancers from Gorkin et al. 2020 supplementary table 8c, or cEnhancers. Wilcoxon rank sum test, * p < 2.2e-16. **D.** Validation rate of indicated elements tested for *in vivo* enhancer activity from the VISTA elements database. Chi-square test, ** p < 0.001, * p < 0.05. **E.** Heatmap for chromatin features and expression of E12.5 cEnhancer-gene pairs. Pairs were k-means clustered by H3K27ac signal surrounding 2 Kb at center of cEnhancer. **F.** Top enriched GO: Biological Process terms for genes from clusters in Fig. 3E. **G.** Example gene, *Neurod1*, showing correlated E12.5 tissue-specific H3K27ac signal at cEnhancer, gene expression, and cEnhancer-gene interactions. The cEnhancer-gene interactions in this region from each E12.5 tissue are marked by arcs. *Neurod1* TSS region was highlighted by yellow box and the cEnhancers are highlighted by blue boxes. **H.** Enriched known and *de novo* motifs that have TF gene expression that matches cEnhancer H3K27ac tissue patterns.

H3K27ac signal at distal enhancers has been shown to correlate with the expression level of their target genes^41, 43^. Assuming a subset of these gene-interacting cCREs are enhancers, we performed the correlation analysis to determine which cCREs likely function as enhancers to their interacting genes. Specifically, we calculated the Spearman’s correlation coefficient between the H3K27ac signal at the distal cCRE with gene expression levels (mRNA-seq signals) of the interacting gene across 22 replicates of tissue/timepoint combinations (**Table S2**, see **Methods** for details). We found the distribution for correlations between H3K27ac signal at cCRE and mRNA-seq from the interacting genes was not only higher than that between randomly shuffled pairs (**Fig. 3B**), but also higher than that between cCREs and their closest promoters (**Fig. S7B**), supporting an active role for 3D chromatin interactions in gene regulation. We classified 16,802 candidate enhancer-target gene pairs (cEnhancer-gene pairs) as those that have a positive correlation (FDR < 0.05) (**Fig. 3B, Fig. S7A**). Of the total 15,098 candidate enhancers identified in these pairs, 90% had one predicted target gene (**Fig. S7C**), while most predicted target genes had multiple assigned enhancers (**Fig. S7D**). Most candidate enhancers were predicted to target promoters more than 200 Kb away (**Fig. S7E**), with 64% of them targeting a gene that is not the nearest promoter (**Fig. S7F**). Further supporting a role for the enhancers predicted to regulate target genes in the mouse embryonic tissues, we found that they had higher sequence conservation than either a random set of size-matched genomic regions or the full list of cCREs (**Fig. 3C**). Importantly, they also had higher sequence conservation than the subset of cCREs (**Fig. 3C**, “Gorkin et al.”) predicted to regulate target genes based on only correlated ChIP-seq/RNA-seq signals without consideration of 3D interaction data^7^. To further assess the validity of these enhancers, we compared them to the elements in the VISTA database^44^. VISTA elements are experimentally tested for enhancer activity by an *in vivo* reporter assay in E11.5 or E12.5 mouse embryos. More than 3,000 elements have been reported to be either positive or negative for enhancer activity in one or more tissues. Referencing the VISTA database, we found that the above enhancers had a significantly higher rate for positive enhancer activity than all cCREs, interacting cCREs, and enhancers predicted without considering 3D interactions (“Gorkin et al.”) (**Fig. 3D**). Taken together, our list of enhancers is supported by multiple lines of evidence for their role in spatiotemporal expression in the mouse embryos. These analyses highlight the power of integrating cCRE-gene activity correlation data with 3D interaction information for enhancer prediction.

Cell-type- and tissue-specific gene expression is often dependent on enhancers for activation^45^, therefore, we hypothesized that genes interacting with predicted candidate enhancers are highly expressed in tissues where they are active. Indeed, genes interacting with VISTA enhancers through the above enhancer-gene links were expressed relatively high in the same tissue where enhancer activity was reported (**Fig S7G**). In several instances, enriched GO terms for genes interacting with VISTA enhancers were related to the tissue where the VISTA enhancer was active. For example, we found that FB positive VISTA enhancers interact with genes expressed the highest in FB and function in forebrain development, while CF positive VISTA enhancers interact with genes expressed the highest in CF and function in cartilage and skeletal system development. These results support a functional relationship between interacting enhancers and promoters and demonstrate the importance of these interactions for tissue-specific activation of gene expression during mouse fetal development.

### Tissue-specific gene regulatory programs during mouse fetal development

To investigate the tissue-specificity gene expression programs in mouse embryos, we focused on the predicted enhancer-gene pairs. We performed K-means clustering of H3K27ac signals at the above enhancers for pairs that had a significant PLAC-seq interaction in one or more E12.5 tissues. These pairs were clustered into 4 main groups (K1-K4) with tissue-specific H3K27ac signals (**Fig. 3E, Fig. S8A**). Candidate enhancers in clusters K1, K2 and K3 were marked with high H3K27ac signals in neural tissues, limb and craniofacial prominence, and liver tissues, respectively, while candidate enhancers in cluster K4 had low H3K27ac signals across all seven tissues. Clusters K1 and K2 were sub-clustered into K1-a, K1-b, K1-c, K2-a, K2-b, K2-c based on the overall H3K27ac signal strength (**Fig. 3E**). We examined additional histone modifications that correlate with activation and enhancer classification (H3K9ac and H3K4me1), as well as the repressive mark H3K27me3 at candidate enhancers. We observed a high degree of correlation between H3K27ac, H3K9ac, and H3K4me1, which had a negative correlation with repressive mark H3K27me3. Similar to H3K27ac, these additional histone marks displayed highly tissue-specific patterns and further support candidate enhancers as putative functional regulatory elements.

While clusters K1-K3 displayed high signal for active histone marks in one or more tissues, K4 displayed low signal for active histone marks in all seven tissues. We observed relatively high CTCF signal across all seven tissues for cluster K4, suggesting that CTCF may be involved in mediating this subset of promoter interactions in the absence of active histone marks (**Fig. 3E**). Interestingly, while clusters K1 and K2 had relatively low but tissue-specific CTCF signal, liver-specific active cEnhancers (K3) had high CTCF binding in all tissues, suggesting that liver-specific developmental enhancers are particularly enriched for CTCF binding (**Fig. 3E**).

To determine the degree of tissue-specificity for long-range chromatin interactions at E12.5 candidate enhancer-gene pairs, we plotted normalized contact frequencies (**Fig. 3E**). Tissue-specific contact strengths strongly correlated with candidate enhancer H3K27ac signal, demonstrating that enhancer activity correlates with the strength of long-range promoter-anchored chromatin interactions. Conversely, there was a strong negative correlation between cEnhancer H3K27me3 signal and contact strength (**Fig. 3E**), suggesting that Polycomb-mediated repression may function to prevent aberrant enhancer-promoter contacts for proper tissue specification during mouse fetal development. Additionally, target genes regulated by E12.5 cEnhancers demonstrated tissue-specific expression and active histone marks at their promoter regions (**Fig. 3E**). Expression levels negatively correlated with H3K27me3 signal (**Fig. 3E**), suggestive of pervasive Polycomb-mediated repression of tissue-specific gene expression by promoter silencing in addition to distal enhancer silencing. Histone modification signals at target promoters generally correlated with signals at candidate enhancers. Taken together, this integrative analysis showcases the high degree of tissue-specific activity of enhancer-promoter pairs during mouse fetal development.

Gene ontology analysis of cEnhancer target genes from E12.5 clusters revealed high enrichment for tissue-specific developmental genes (**Fig. 3F**). For example, genes targeted by cEnhancers active in neural tissues (K1) had the highest enrichment for terms “nervous system development” (p = 1×10^-255^) and “neurogenesis” (p = 1×10^-205^). Genes targeted by cluster K2 cEnhancers active in limb and craniofacial prominence were related to anatomical structure morphogenesis (p = 1×10^-196^), while genes targeted by liver-active cEnhancers function in cellular macromolecule metabolic process (p = 1×10^-96^). Cluster K4, which had no apparent tissue-specific features at cEnhancers, had genes most enriched for the term “system development” (p = 1×10^-152^) indicating that these genes are less specific for development of any particular tissues. These data demonstrated the high degree of tissue-specificity of long-range enhancer-promoter interactions for tissue-determining gene expression during mouse fetal development. For example, *Neurod1*, a brain-specific TF involved in neuron differentiation, was selectively expressed in nervous system and was assigned to distal cEnhancers only in FB, MB, HB, and NT (**Fig. 3G**).

To identify potential developmental regulators involved in establishing candidate enhancers, we performed TF-motif enrichment analysis for these regions. We found highly enriched TF-motifs for each cluster and filtered the top enriched TFs for those with tissue expression patterns that matched with cEnhancer activity (**Fig. 3H**). This analysis produced a high confidence list of TFs that likely activate tissue-specific developmental enhancers (**Fig. 3H**). Within this list many known developmental regulators were identified in the expected clusters. For example, K1 was enriched for *Rfx*-family, *Sox*-family, *Pou3f3*, *Neurod1*, and *Ascl1* TF motifs, all of which have been previously reported to play important roles in nervous system development^46-50^. *Foxo1* and *Atoh8*, important for chondrogenic commitment of skeletal progenitor cells^51, 52^, and *Myog* and *Six1*, important for skeletal muscle fiber differentiation^53^, were enriched at K2 cEnhancers active in limb and craniofacial prominence. Liver-specific cEnhancers had enrichment for hepatocyte differentiation factors *Gata4* and *Cebpb*^54^. Finally, we found that the CTCF motif had the highest enrichment for K4 cEnhancers, supporting the high CTCF binding detected by ChIP-seq at these cEnhancers (**Fig. 3E**). Our results implicated several transcription factors to bind distal enhancers that physically interact with target promoters and activate tissue-specific gene expression.

### Dynamic enhancer-promoter interactions in developing forebrain

We also performed K-means clustering of H3K27ac signal at the cEnhancers from cEnhancer-gene pairs with a significant PLAC-seq interaction present in forebrain between E12.5 and P0, and produced 5 clusters (**Fig. S8A**), with K1, K2, and K3 showing highly dynamic changes in active and repressive histone marks through developmental stages (**Fig. S8B**). Interestingly, the chromatin interaction strengths were also dynamic across developmental stages in a pattern similar to H3K27ac. Consistent with E12.5 clusters (**Fig. 3E**), active histone marks at promoters correlated with those at interacting cEnhancers though with less overall dynamics (**Fig. S8B**). We then integrated previously generated single-nucleus ATAC-seq data from the same fetal mouse forebrain samples^55^ to identify specific forebrain cell types contributing to the cEnhancer activity from these clusters (**Fig. S8B**). Cluster K1 cEnhancers gained signal for active histone marks as development progressed, which appeared to be contributed mostly by eEX2, excitatory neurons. Excitatory neurons have low abundance at stage E12.5 but become more abundant as development progresses. In particular, eEX2 cells are the most abundant cell type at P0^55^, suggesting that eEX2 abundance contributes to these tissue-wide H3K27ac dynamics at K1 cEnhancers. Conversely, H3K27ac at K2 cEnhancers gradually decreased from E12.5 to P0, which reflects these sites having closed, inactive chromatin in eEX2 cells (**Fig. S8B**). The integration of cell-type-specific open chromatin data demonstrates that forebrain enhancer dynamics during fetal development is correlated with changes in cellular composition.

We performed GO term analysis on genes from each forebrain candidate enhancer-gene pair cluster. For clusters K1-K4 the most enriched term was nervous system development (**Fig. S8C**) while cluster K5, which had low H3K27ac signal and high CTCF binding at candidate enhancers at all developmental stages (**Fig. S8B**), displayed GO enriched terms related to development without specific tissue designation (**Fig. S8C**). An example of a gene with temporally dynamic enhancer-promoter interactions is *Neurod6*, a TF important for neuronal differentiation^56^. Expression of *Neurod6* is low at E12.5, where there are no assigned interacting cEnhancers, but becomes activated at E13.5 coinciding with the onset of several cEnhancer interactions at H3K27 acetylated distal sites (**Fig. S8D**). These distal interacting cEnhancers are marked with open chromatin in excitatory neurons, cells known for high *Neurod6* expression^57^. TF motifs enriched at cEnhancers with TF expression patterns that match cEnhancer activity implicated many known brain developmental regulators as activators for these enhancers (**Fig. S8E**). For each cluster, this revealed unique TFs likely contributing to the diverse temporal activity of these cEnhancers during forebrain development. For example, cEnhancers active at E12.5 are likely activated by TFs such as *Lhx2* and *Ascl1*, while cEnhancers more active at later stages are likely established by different TFs such as *Neurod6* and *Mef2*. Our results demonstrate that extensive temporal dynamics of enhancer-promoter interactions regulate the precision of stage-specific gene expression during mouse fetal brain development.

### Enhancer-promoter interactions facilitate interpretation of noncoding risk variants in the human genome

Genome-wide association studies (GWAS) have identified hundreds of thousands of common genetic variants linked to various traits and diseases. However, most GWAS variants reside in non-coding regions, and it is still very challenging to interpret their functions^58^. To address this challenge, we utilized our list of candidate enhancer-gene pairs in the mouse genome to predict gene targets for human enhancers harboring non-coding GWAS variants. We first identified 12,929 human orthologous regions corresponding to the 15,098 mouse candidate enhancers, of which 68% overlap human cCREs in the ENCODE registry (https://screen.encodeproject.org). Using these human orthologous regions, we performed linkage disequilibrium score regression (LDSC) on a panel of 65 GWAS phenotypes. Clusters of E12.5 active candidate enhancers (**Fig. 3E**) were enriched for GWAS variants associated with specific phenotypes (**Fig. 4A**). In most cases, enrichments displayed tissue-specificity that matched with tissues expected to contribute to the associated phenotype (**Fig. 4A**). For example, neural-active K1 candidate enhancers were found enriched for non-coding variants associated with neurological disorders such as depression, autism, schizophrenia, and neuroticism. K2 enhancers active in limb and craniofacial prominence were enriched for non-coding variants associated with physical size, and K3 enhancers active in liver had the highest enrichment for non-coding variants associated with metabolic diseases. The enrichment of GWAS variants at tissue-specific fetal mouse enhancers indicates the potential of our annotated mouse enhancer-gene pairs for understanding mechanisms related to disease caused by non-coding genetic variants.

**Figure 4.**
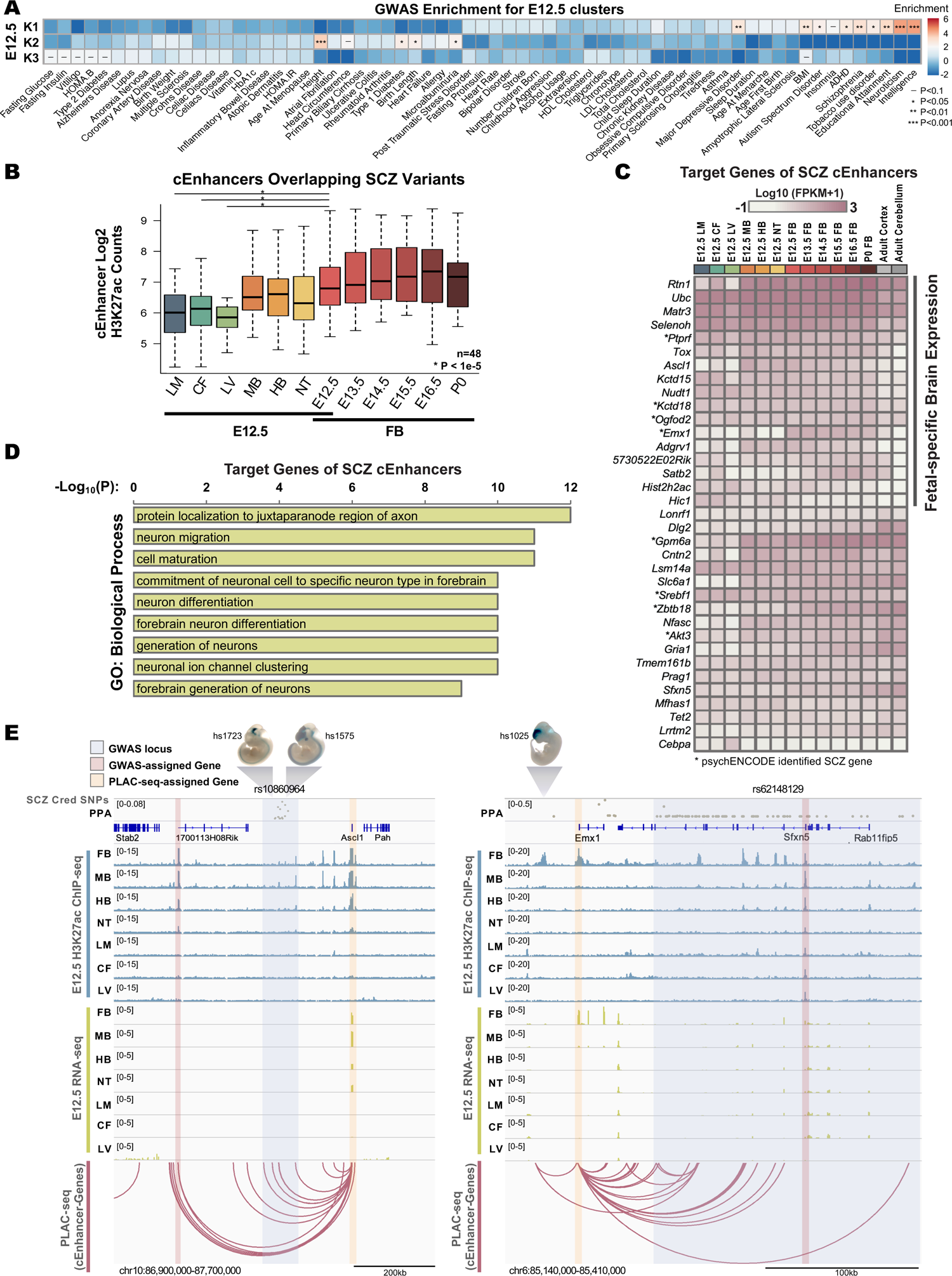
Chromatin interactions facilitate interpretation of noncoding disease risk variations in the human genome. **A.** Linkage disequilibrium score regression analysis to identify GWAS enrichments at cEnhancers from E12.5 clusters shown in Fig. 3E. **B.** Box plots showing distribution of H3K27ac counts at cEnhancers overlapping SCZ credible set SNPs. Two-tailed paired T-Test between E12.5 FB and non-neural tissues. **C.** Heatmap of gene expression for genes identified as interacting with cEnhancers overlapping SCZ credible set SNPs, n=35, psychENCODE SCZ genes^50^. Adult mouse cortex and cerebellum RNA-seq data obtained from ENCODE. **D.** Gene ontology analysis showing top enriched Biological Process terms of genes interacting with cEnhancers harboring SCZ credible set SNPs. **E.** SCZ example genes, *Ascl1* and *Emx1*, interacting with loci harboring SCZ-candidate SNPs. Tracks display H3K27ac, gene expression, and all identified nearby cEnhancer-gene interactions. Loci are labelled with Tag SNPs rs10860964 and rs62148129. Select images of mouse embryos displaying positive VISTA enhancer activity in the displayed genomic regions were obtained from VISTA database. All cEnhancer-gene pairs with a PLAC-seq interaction in any tissue/stage tested present in these genomic regions are marked by arcs.

Schizophrenia (SCZ) is a neurological condition in adulthood that is associated with many genetic variants, however, the disease etiology remains poorly understood^59^. To predict putative target genes of SCZ-associated non-coding variants, we focused on a 99% credible set of 20,373 SCZ SNPs^60^ (obtained from the Psychiatric Genomics Consortium website at http://www.med.unc.edu/pgc/) concentrated at 266 genomic loci. We identified 48 human orthologous candidate enhancers containing SCZ-associated SNPs that target 35 genes (**Fig. 4B, Fig. 4C**). The mouse enhancers had high H3K27ac signal in neural tissues (**Fig. 4B**) and their predicted target genes display high levels of gene expression (**Fig. 4C**). The predicted target genes were highly enriched for GO terms related to neuronal differentiation (**Fig. 4D**), and included transcription factors such as *Ascl1*, *Emx1*, *Tox*, and *Zbtb18,* known to regulate neuronal developmental (**Fig. 4C**). However, genes closest to these 48 mouse candidate Enhancers lacked brain-specific GO term enrichment (**Fig. S9A**). Interestingly, 17 of the predicted target genes are expressed in fetal mouse brain but become repressed in adult brain (**Fig. 4C**), suggesting that aberrant gene expression during fetal brain development may contribute to SCZ, a disease with onset most common in early adulthood. Further, many of our predicted target genes were not identified in studies using non-fetal human data^61^, demonstrating that fetal mouse enhancer-promoter interactions add additional insights into understanding molecular mechanisms contributing to SCZ.

For example, *Ascl1*, a regulator of neuroblast differentiation, forms long-range chromatin interactions in fetal mouse forebrains with candidate enhancers, human orthologous locus of which harbor SCZ SNPs. By proximity the SCZ risk variants are assigned to the nearest and uncharacterized gene *C12orf42*, however, chromatin interactions data from the mouse forebrain revealed that the enhancers harboring these risk variants likely regulate *Ascl1* (**Fig. 4E**). Two VISTA elements confirm this locus exhibits brain and neural tube-specific enhancer activity during fetal development (**Fig. 4E**). Due to its fetal-specific enhancers and gene expression in brain (**Fig. S9B**), *Ascl1* was not identified previously as a candidate SCZ causal gene^61^. Studies have consistently found abnormalities in thalamocortical axonal connectivity in SCZ patients^62^, and since it is critical for relaying sensory information from the periphery to the neocortex, thalamocortical network abnormalities in SCZ have been postulated to contribute to hallucination events^63^. Supporting the assignment of *Ascl1* as a target gene of the SCZ risk variants, *Ascl1* null mice exhibited defects in thalamocortical axonal connectivity^64^, providing a physiological link between *Ascl1* disruption and neurocognitive abnormalities in SCZ. Additionally, *Emx1*, a regulator of neuronal commitment, had high expression in fetal forebrain, but was repressed in adult brain (**Fig. 4C, Fig. S9B**). We identified *Emx1* promoter interactions with several candidate enhancers marked with forebrain-specific H3K27ac including VISTA element hs1025, which displayed fetal enhancer activity exclusively in the forebrain (**Fig. 4E**). The human orthologous sequences of the *Emx1*-interacting candidate enhancers harbor SCZ-risk SNPs, however, by proximity, these risk variants are assigned to a different gene, *Sfxn5* (**Fig. 4E**), a widely expressed mitochondrial tricarboxylate carrier^65^. These examples showcase the value of our catalogue of developmental enhancer-gene pair predictions for molecular understanding of how genetic variation contributes to complex human disease.

## Discussion

In this study, we comprehensively profile the promoter-centered chromatin interactomes in twelve mouse fetal tissues using H3K4me3 PLAC-seq and identified a total of 248,620 long-range chromatin interactions. In accordance with previous reports, the long-range interactions exhibit both tissue-to-tissue variability and developmental dynamics, and the promoter-interacting regions are enriched for tissue-specific open chromatins and enhancer marks. Integrating this dataset with CTCF ChIP-seq data also reveals a prevalent involvement of CTCF in facilitating promoter-enhancer contacts in various fetal tissues.

Annotating target genes of candidate enhancers is a major challenge confronting geneticists today. First, accurately defining functional enhancers from epigenomic datasets alone is nontrivial: our previous study found that only ∼30-60% of putative enhancers with known enhancer marks (such as H3K27ac signal) drive reporter genes in corresponding tissues *in vivo* in transgenic assays^7^. Second, deletion of an individual enhancer often does not significantly alter the expression level of its target gene assigned by physical contacts, probably because the deleted enhancer is redundant, or the detected enhancer-promoter contact is too weak or simply false positive signal. To improve the accuracy in predicting functional enhancer-promoter pairs, we took advantage of the rich datasets from 12 mouse fetal tissues and integrated 3D chromatin interaction datasets with epigenomic and transcriptomic datasets for such prediction. We assume that the functional enhancer-promoter pairs not only show significantly enriched physical contacts from our PLAC-seq data but also have positive correlation on their epigenomic signals/gene expression across different tissue/cell types. As a result, we found in total 16,802 candidate enhancer-gene pairs and further leveraged these pairs to assign non-coding genetic variants associated with various human diseases and traits. Specifically, the analysis on SCZ-associated non-coding variants revealed potential SCZ-associated genes that exclusively expressed in fetal mouse brain that were not identified in previous studies using non-fetal human data. Such analysis showcases the value of utilizing mouse fetal omic data to reveal novel insights into human disease etiology.

Despite the comprehensive datasets included in this study, there are still a few important limitations. First, the prediction of candidate enhancer-gene pairs is solely relied on the omics data while functional validations, such as CRISPR depletion of candidate enhancers in relevant cell types or tissues followed by gene expression profiling, are necessary to experimentally validate the contribution of candidate enhancers to the target gene expression. Second, all the tissues used in this study are heterogeneous. Epigenome and 3D genome information at single cell resolution is critical to further delineate the specific cell types contributing to the changes in candidate enhancer activity and long-range interactions. For example, the activity change on different groups of candidate enhancers in forebrains is likely to be caused by the cell composition changes during development, based on the single-nucleus ATAC-seq data^55^ from the same samples (**Fig. S7B**). Finally, not all the fetal tissues studied in the ENCODE project Phase III were included in this study due to exhaustion of tissue samples. Nevertheless, we believe that the findings presented here demonstrate the potential of integrating multi-omics datasets to discover new mechanistic insights of gene regulation. Such insights gained from early mouse fetal tissues may also be extrapolated to infer gene regulatory networks influencing human development and disease.

## Data availability

The H3K4me3 PLAC-seq and CTCF ChIP-seq data have been deposited to GEO with accession number GSE200114 and available from 4DN Data portal (https://data.4dnucleome.org).

## Acknowledgements

This study was funded by NIH grants U54DK107977 (B.R.), UM1HG011585 (B.R.) and R35HG011922 (M.H.).

## Author Contributions

This study was conceived and designed by M.Y., M.H. and B.R.; Experiments were performed by M.Y. and R.H.; Data analysis was performed by M.H., N.Z., Z.C., I. J., M.Y., R.R., A.A. and R.F.; Manuscript was written by M.H., M.Y. N.Z., Z.C. and B.R. with input from all authors.

## Methods

### Tissue collection

All mouse tissues used in this study are collected for the ENCODE project and the step-by-step protocol can be found on the ENCODE project website: https://www.encodeproject.org/documents/631aa21c-8e48-467e-8cac-d40c875b3913/@@download/attachment/StandardTissueExcisionProtocol_02132017.pdf.

### Tissue fixation

30-50 mg tissues are crosslinked with 1% formaldehyde (v/v) in 1XPBS buffer with 0.1 M NaCl, 1 mM EDTA, 0.5 mM EGTA and 50 mM HEPES (pH 8.0) at room temperature for 15 min with slow rotation. The fixation was quenched by addition of 2.5 M glycine solution to a final concentration of 0.125 M with slow rotation at room temperature for 5 min. Fixed tissues were pelleted by centrifugation at 2,000×*g* for 5 min at 4 °C and washed with ice-cold PBS once. The washed tissues were pelleted again by centrifugation, snap-frozen in liquid nitrogen and stored at −80 °C.

### H3K4me3 PLAC-seq data generation

30-50mg frozen tissues were thawed on ice and then resuspended in 1ml of lysis buffer (10 mM Tris-HCl, pH 8.0, 5 mM CaCl_2_, 3 mM MgAc, 2 mM EDTA, 0.5 mM EGTA, 1 mM DTT, 0.1 mM PMSF with proteinase inhibitor). Tissue dissociation was performed using the gentleMACS™ dissociator with program “Protein-M-tube-1.0”. After dissociation the sample was filtered with 40 µm cell strainer to remove large particles and additional lysis buffer with 0.4% Triton X-100 was added at equal volume to make the final concentration of Triton X-100 as of 0.2%. The nuclei were centrifuged at 1,000×*g* for 6 min at 4°C. The supernatant was discarded and 3 ml sucrose buffer (1M sucrose, 10 mM Tris-HCl, pH 8.0, 3 mM MgAc with proteinase inhibitor) was carefully added from the side of the tube. The nuclei were centrifuged at 2,500×*g* for 6 min at 4 °C and the supernatant was discarded. The nuclei are ready for the PLAC-seq experiment. PLAC-seq libraries were prepared as described in previous study^16^ with minor modifications and the detail protocol is available via 4DN portal: https://data.4dnucleome.org/protocols/4ba366ad-b261-4545-baa0-89776c3ab699/.

### CTCF ChIP-seq data generation

∼30 mg fixed tissues were thawed on ice and the nuclei preparation was performed using truChIP® Chromatin Shearing Kit (Covaris) following the manufacturer’s instruction. The lysed nuclei were sheared using Covaris M220 with following setting: Setting: power, 75 W; duty factor, 8%; cycle per burst, 200; time, 12 min; temp, 7. The sheared chromatins were cleared by centrifugation at 15,000×*g* for 12 min. The supernatant was collected and incubated with M-280 sheep anti-rabbit magnetic beads (Thermo Fisher, 11203D) for 3 hours at 4°C with slow rotation for preclearing. ∼5% of precleared cell lysate was saved as input control after the incubation was done. The rest of precleared lysate was mixed M-280 sheep anti-rabbit magnetic beads that coupled with 0.9 μg of CTCF antibody (3418, Cell Signaling) and rotate at 4°C for 16 hours. On the next day, the beads were collected using magnetic stand and the supernatant was discarded. The beads are then washed with RIPA buffer three times, high-salt RIPA buffer (10 mM Tris, pH 8.0, 300 mM NaCl, 1 mM EDTA, 1% Triton X-100, 0.1% SDS, 0.1% sodium deoxycholate) twice, LiCl buffer (10 mM Tris, pH 8.0, 250 mM LiCl, 1 mM EDTA, 0.5% IGEPAL CA-630, 0.1% sodium deoxycholate) once, TE buffer (10 mM Tris, pH 8.0, 0.1 mM EDTA) twice. Washed beads were treated with 10 μg Rnase A in extraction buffer (10 mM Tris, pH 8.0, 350 mM NaCl, 0.1 mM EDTA, 1% SDS) for 1 hours at 37°C, followed by reverse crosslinking in the presence of proteinase K (20 μg) overnight at 65°C. After reverse crosslinking the DNA was purified and quantified. The input control DNA was reverse crosslinked and purified in the same way as ChIPed DNA as described above. Library preparation was performed using QIAseq Ultralow Input Library Kit (Qiagen) starting from 10–100 ng ChIPed DNA or input DNA. After PCR the amplified libraries were purified with SPRI Beads to extract fragments between 200-600bp for sequencing.

### CTCF ChIP-seq data processing and peak calling

The fastq files of CTCF ChIP-seq data were mapped to mouse genome (mm10) and processed using the ENCODE uniform processing pipeline for ChIP-seq data (https://github.com/ENCODE-DCC/chip-seq-pipeline) with default parameters.

Peaks were called using MACS2^66^ with regular peak calling at *P* threshold of 0.01. Such relaxed peak sets were generated for each biological/technical replicate, and also for the pooled replicates. Peaks from the pooled replicate set were defined as the replicated peak set if they overlapped (at least 1 bp) the peaks from both biological/technical replicates.

### H3K4me3 PLAC-seq data processing and interaction calling

We applied the MAPS pipeline (https://github.com/HuMingLab/MAPS) to analyze H3K4me3 PLAC-seq data, as described in our previous study^16^. Briefly, we first mapped the raw fastq files to the reference genome mm10 and only kept the uniquely mapped reads. We then selected valid read pairs after removing PCR duplicates. The deduped, valid read pairs were used to generate the raw contact matrix (without normalization) in the .hic format using Juicer^67^ for visualization.

We then partitioned the genome into 10 Kb bins, and counted the number of paired-end reads in each 10 Kb bin pair. Next, we defined the “AND”, “XOR” and “NOT” sets as 10 Kb bin pairs where both two ends, only one end, and neither end contain replicated H3K4me3 ChIP-seq peaks (the “replicated peaks” files from Encode: https://www.encodeproject.org, see also **Table S3**) from the corresponding tissue, respectively. We only kept the bin pairs in “AND” and “XOR” sets with >=1 counts for the downstream analysis of interaction calling. We assumed that the counts in the “AND” and “XOR” sets follow the positive Poisson distribution, and fitted a positive Poisson regression to adjust for systematic biases from effective fragment size, GC content, mappability, ChIP enrichment level (measured by the number of short-range [<1 Kb] reads within each 10 Kb bin), and 1D genomic distance. Next, we defined the normalized contact frequency as the ratio between the observed contact frequency and the expected contact frequency obtained from the positive Poisson regression. Finally, we identified long-range significant chromatin interactions at 10 Kb resolution in autosomal chromosomes between 1D genomic distance range of 20 Kb ∼ 1 Mb, such that these 10 Kb bin pairs satisfied: (1) FDR<1% and (2) normalized contact frequency >=2. The significant interactions in the “AND” set are referred as “Peak-to-Peak” interactions and the ones in the “XOR” set are referred as “Peak-to-Non-peak” interactions.

### Quality control of PLAC-seq data

The “feather” part of MAPS is a preprocessing pipeline tailored to PLAC-Seq/HiChIP data and it outputs a “.feather.qc” file that contains metrics to evaluate the quality of PLAC-Seq library. Among these metrics, the most important ones are “trans_ratio”, “long_cis_ratio” and “FRiP (Fraction of Reads in Peaks)” values. We require all replicates in our study to satisfy the following requirements.

1) trans_ratio <40%: trans_ratio is calculated as the number of inter-chromosomal read pairs in the final bam file divided by the number of uniquely mapped read pairs that were kept after PCR duplication removal.

2) >50% long_cis_ratio: long_cis_ratio is calculated as the number of long-range (defined by the input argument to feather, default >1,000 bp) intra-chromosomal read pairs in the final bam file after removing alternative and mtDNA chromosomes (if any), divided by the number of all intra-chromosomal read pairs in the final bam file after removing alternative and mtDNA chromosomes (if any).

3) >7.5% FRiP: FRiP is calculated as the number of “valid” short-range reads (the two ends of reads coming from two different strands, intra-chromosomal, <=1,000bp) in the final bam file after removing alternative and mtDNA chromosomes (if any) reads that overlap with the reproducible H3K4me3 ChIP-seq peaks in the same tissue, divided by the total number of “valid” short-range reads.

### Reproducibility analysis of MAPS-identified interactions from two biological replicates

After acquiring the MAPS-identified long-range chromatin interactions from each biological replicate, two approaches were used to evaluate their reproducibility: (1) we calculated the overlap between MAPS-identified interactions from two replicates, (2) we calculated the Pearson correlation coefficients of the normalized contact frequency for the reproducible interactions from two replicates.

### Merging and down-sampling of H3K4me3 PLAC-seq data from the two biological replicates

The PLAC-seq data from the two biological replicates were combined by summing up the read counts in 10 Kb bin pairs in the “AND” and “XOR” sets. To account for the variable sequencing depths, we further down-sampled above combined data. Specifically, for each of 19 autosomal chromosomes, we counted the total number of paired-end reads in the “AND” and “XOR” sets across 12 tissues. We identified the tissue with the minimal number of reads, and randomly down-sampled the paired-end reads in the “AND” and “XOR” sets in the remaining 11 tissues to match that minimal number. After such down-sampling procedure, we ensured that for each chromosome, the total number of paired-end reads in the “AND” and “XOR” sets are the same across 12 tissues.

### Interaction calling from the combined, down-sampled data

MAPS was applied to the combined, down-sampled data of each tissue as described in the “**H3K4me3 PLAC-seq data processing and interaction calling**” section to identify long-range interactions.

### Normalized contact frequency calculation and interaction calling using union H3K4me3 peaks list

Since only the AND and XOR sets were taken into consideration in MAPS, a bin pair identified as significant interaction from one tissue might not be testable in another tissue. Thus, for valid comparison across 12 tissues, we re-defined the AND and XOR sets of 10 Kb bin pairs in each tissue by taking the union of H3K4me3 ChIP-seq peaks from all 12 tissues and re-calculate the normalized contact frequency and long-range interactions as described above in the “**H3K4me3 PLAC-seq data processing and interaction calling**” section.

In this paper, for comparison analysis across tissues (Figure 2, Figure S5 and Figure S6), we used normalized contact frequency and MAPS interaction identified using union H3K4me3 ChIP-seq peak list. Otherwise, we used replicated H3K4me3 ChIP-seq peaks from the corresponding tissue for MAPS.

### Gene and TSS Annotation

We downloaded Gencode vM4 annotation (https://www.encodeproject.org/data-standards/reference-sequences/) to define transcript TSS for each protein coding gene. To count number of genes involved in or in PLAC-seq interactions, we only considered whether the TSS of the gene is laid in either of the 10 Kb ends of the PLAC-seq interactions.

### Comparison of MAPS-identified interactions with Capture-C interactions

The Capture-C interactions from forelimb and hindlimb at E10.5, E11.5 and E13.5, and midbrain at E12.5 were downloaded from Gene Expression Omnibus (GEO) with accession number GSE84795, using the 95^th^ or the 99^th^ percentile as the cut-off value^27^. The mm9 coordinates used in the original paper were converted to mm10 using UCSC LiftOver tool (https://genome.ucsc.edu/cgi-bin/hgLiftOver). Since MAPS only detected interactions that have at least one end overlapping H3K4me3 peaks, we further filtered Capture-C interactions for fair comparison. Specifically, a Capture-C interaction was considered as testable only when it satisfies the following two criteria: (1) the interaction was with 1D genomic distance between 20 Kb to 1 Mb in autosomal chromosomes and (2) the interaction has at least one end overlapping the 10 Kb bins containing reproducible H3K4me3 peaks identified in the corresponding tissue (LM E12.5 for forelimb and hindlimb Capture-C data and MB E12.5 for midbrain E10.5 Capture-C data).

Since PLAC-seq and Capture-C were not performed on exactly the same tissues, the MAPS-identified interactions of MB E12.5 were compared with Capture-C interactions from midbrain E10.5, and the MAPS-identified interactions of LM E12.5 were compared with Capture-C interactions from forelimb and hindlimb. A testable Capture-C interaction was considered as being recapitulated by PLAC-seq when its both ends overlapped the two ends (at least 1 bp) of a PLAC-seq interaction identified from the corresponding tissues. As a control set, a list of distance-matched pseudo-interactions was generated by linking the H3K4me3-containing 10 Kb bin to the 10 Kb bin equidistant from but on the other side of H3K4me3-containing bins.

### Enrichment analysis of open chromatins and histone marks in promoter-interacting regions

For Promoter-to-Non-promoter interactions, we defined the 10 Kb bins which did not contain H3K4me3 peaks as promoter-interacting regions and asked whether they are enriched for open chromatins and/or specific histone modifications compared to distance-matched control (the 10 Kb bin equidistant from but on the other side of H3K4me3-containing bins). The replicated peaks of each histone modification and the replicated ATAC-seq peaks of each tissue were downloaded from ENCODE data portal (https://www.encodeproject.org/) and the identifiers are summarized in **Table S3**. Enrichment score were defined as the fold-change of the proportion of promoter-interacting regions and the control regions overlapping the midpoint of the replicated histone ChIP-seq peaks or ATAC-seq peaks from the same tissues.

### Enrichment analysis of chromatin states on promoter-interacting regions

The ChromHMM annotation files of all samples used in this study were downloaded from http://enhancer.sdsc.edu/enhancer_export/ENCODE/chromHMM/replicated/ (replicated)^7^. The promoter-interacting regions and the control regions, which have been defined in the “**Enrichment analysis of open chromatins and histone marks in promoter-interacting regions”** section, were overlapped with the reproducible autosomal chromHMM state calls from the same tissue. Enrichment score (fold-change) of a given state “S” in a particular tissue was calculated as the number of base pairs of the promoter-interacting regions that overlap state “S”, divided by the number of base pairs of the control regions that overlap state “S”.

### Enrichment analysis of CTCF on interaction anchors

To calculate the expected proportion of such interactions with at least one end bound by CTCF for P2P interactions, we first checked the co-occupancy of peaks identified from CTCF ChIP-seq and peaks identified from H3K4me3 ChIP-seq across whole genome, estimating *p* - the probability for a randomly selected H3K4me3-containig 10 Kb bin to have CTCF binding. Then, we calculated the expected proportion for a P2P interaction that has CTCF binding on at least one end as 1 − (1 − *p*)^2^. Since the CTCF peaks identified from ChIP-seq data varied from sample to sample, the expected proportion was calculated for each sample respectively and we used paired t-test to calculate the p-value by comparing the observed fraction and expected fraction in 12 samples (seven tissues from E12.5 and five extra time points of forebrain).

Similarly, for Promoter-Non-promoter (P2N) interactions, we also calculated *q* – the probability for a H3K4me3-depleted 10-bin to have CTCF binding. Then, the expected proportion for a P2N interaction that has CTCF binding on at least one end was calculated as 1 − (1 − *p*)(1 − *q*) and then paired t-test was applied on the same 12 samples.

### CTCF motif orientation analysis

FIMO^68^ with default parameters was used to search for all CTCF motifs (MA0139.1 from the JASPAR database)^69^ within the replicated peaks called from CTCF ChIP-seq data. If more than one CTCF motifs were found in one peak region, we only kept the one with the highest FIMO score. We then intersected the above list with the 10 Kb MAPS-identified interactions and calculated the proportion of convergent, forward-forward, reverse-reverse and divergent CTCF motif pairs among all interactions that have CTCF binding on both ends. It is notable that for some interactions their anchors (10 Kb bin) may contain multiple CTCF peaks with opposite motif orientations. Therefore, we only considered a subset of MAPS-identified interactions with both ends containing either single CTCF motif or multiple CTCF motifs in the same direction.

### Characterization of MAPS-identified interactions by the number of CTCF peaks in the bin

We first classified all H3K4me3-containing 10 Kb bins according to the number of reproducible CTCF ChIP-seq peaks from the same tissue. For the sake of conservation, only CTCF peak with CTCF binding motif in it were considered. Then, we counted number of the Peak-to-Peak (P2P) and Peak-to-Non-peak (P2N) interactions formed on the bins respectively or calculated the fraction of bins forming P2P or P2N interactions in each tissue.

### Interaction direction preference analysis

We first classified all Peak-to-Non-peak interactions which have CTCF binding only on P-end (the end containing H3K4me3 peaks) by asking whether the motif orientation of CTCF binding on the P-end face on an interaction face the direction of the other end of the interaction or is back to the direction of the other end. Then we calculated the proportion of the interactions from two groups.

To estimate the expected value of this proportion, we first selected all 10 Kb bins containing H3K4me3 peaks that were bound by CTCF and then randomly shuffled the CTCF binding, which means, during this process, a CTCF binding on the bin A will be assigned to another bin B, keeping its orientation unchanged. Next, the proportion was recalculated based on the same Peak-to-Non-peak interactions set with shuffled CTCF binding information and this was used as the expected value. Since the CTCF peaks identified from ChIP-seq data varied from sample to sample, the expected proportion was calculated for each sample respectively and paired t-test was applied to calculate the p-value by comparing the observed fraction and expected fraction in 12 samples (seven tissues from E12.5 and five extra time points of forebrain).

### Clustering of PLAC-seq data across 12 tissues

Clustering analysis including principal component analysis (PCA) and hierarchical clustering analysis (HCA) was performed on the normalized contact frequency of 10 Kb bin pairs which were called as significant interactions in at least 1 of the 12 tissues using union H3K4me3 peak list. Bin pairs with zero variance of normalized contact frequency across 12 tissues were discarded. Finally, a total of 195009 bin pairs were used for clustering analysis.

Based on the normalized contact frequency matrix of these 195009 bin pairs, PCA was performed by R function “prcomp” and HCA was performed by the “hclust” function in R using Ward’s minimum variance method (using ‘ward.D’ argument).

### Clustering of H3K27ac ChIP-seq data across 12 tissues

Hierarchical clustering of H3K27ac histone modification ChIP-seq was performed using the “hclust” function in R as previously described^7^. First, replicated H3K27ac ChIP-seq peaks from all 12 tissues were pooled together, merging by bedtools merge (v2.29.2)^70^ so that the same peak list was used across all 12 tissues for comparison. Average fold enrichments on these merged peaks were re-calculated by bigWigAverageOverBed (https://github.com/ENCODE-DCC/kentUtils/blob/master/bin/linux.x86_64/bigWigAverageOverBed), using pooling signal (fold change over control) bigwig files of each tissue as input. Finally, quantile normalization was applied and the normalized scores were used for hierarchical clustering. The replicated peaks of H3K27ac histone modification as well as the signal bigwig files were downloaded from ENCODE data portal (https://www.encodeproject.org/) and the identifiers are summarized in **Table S3**.

### Heatmap for tissue- or stage-specific interaction features

We determined tissue or stage-specific XOR MAPS interactions as those having significance in only the indicated E12.5 tissues (Fig. 2B) or FB developmental stage (**Fig. S6A**). For valid comparisons across samples, we used normalized contact frequencies from the union set anchor interactions. Bedtools^70^ pairToBed function was used to find H3K4me3 marked promoters in the anchor bin and H3K27ac peaks in the non-anchor bin. Heatmap displays the percentage of an individual tissue FPKM of the total sum of FPKM for all tissues for each gene. H3K27ac signal was calculated by counts within peaks overlapping with the non-anchor interacting bins. H3K27ac counts were corrected for total mapped reads and quantile normalized. Heatmap displays the percentage of an individual tissue H3K27ac counts of the total sum of H3K27ac counts for all tissues for each non-anchor bin.

### Correlation analysis between fold-change of gene expression and that of number of MAPS-identified significant chromatin interactions

We first selected 9,939 protein-coding genes which are expressed in at least one of 12 tissues (defined by FPKM>1), and have transcription start site (TSS) overlaps with H3K4me3 ChIP-seq peak in at least one of 12 tissues. Next, we converted the FPKM value to Log2(FPKM+1), and applied R function “normalize.quantile” in the “preprocessCore” library to perform quantile normalization across all 12 tissues. The log2 transformed and quantile normalized gene expression data for these 9,939 genes were used for the downstream analysis. In addition, we counted the number of the number of MAPS-identified significant chromatin interactions in each tissue. For any two tissues, we further made the scatter plot between the change of chromatin interaction and the change of gene expression, and reported the Pearson correlation coefficients and the corresponding P-values.

### Visualization of H3K27ac ChIP-seq and RNA-seq data from mouse fetal tissues

We downloaded the signal tracks from the ENCODE portal (https://www.encodeproject.org/)^71,72^ and the identifiers are summarized in **Table S3**.

### Identification of interacting cCRE-genes and cEnhancer-gene pairs

PLAC-seq interactions between promoter distal elements and active TSSs (XOR set from the MAPS software) were utilized for determining all cCRE-gene interactions. Bedtools intersect (https://bedtools.readthedocs.io/en/latest/) was used to determine all chromatin accessible peaks from the d-TAC catalogue^7^ overlapping a 10 Kb non-anchor bin (cCREs). cCREs were assigned to protein-coding genes with an H3K4me3-marked TSS (Gencode vM4) in the 10 Kb anchor bin for each interaction. This generated a catalogue of 91,451 interacting cCRE-gene pairs. All integrated data were from the same tissue type and at the same developmental stage. To determine candidate enhancers (cEnhancers), these pairs were filtered for having cCRE H3K27ac signal that correlated with interacting gene expression. HTSeq software (https://htseq.readthedocs.io/en/master/) was used to calculate all H3K27ac counts within a 2 Kb window from the center of each interacting cCRE for each tissue and developmental stage. H3K27ac counts were normalized for total counts and values were quantile normalized. Spearman’s correlation coefficients (SCC) for normalized H3K27ac counts at cCREs with the interacting gene’s FPKM value^24^ for each interacting cCRE-gene pair were calculated. Biological replicate datasets for each sample were kept separate unless H3K27ac ChIP-seq and mRNA-seq were generated from different samples, in which case duplicates were combined and averaged. This resulted in 22 data points for SCC calculation (**Table S2**). Those with a positive SCC and FDR < 0.05 were included in the list of cEnhancer-gene pairs. FDR values were calculated from P-values using Benjamini-Hochberg procedure.

### Comparison of the closest promoter cCRE assignment with interacting cCRE promoter pairs

Correlations between H3K27ac at cCREs and expression of assigned target gene were compared between interacting cCRE-gene pairs (n=91,451, described above) and cCRE-closest promoter pairs (n=115,711). For cCRE-closest promoter pairs, only cCREs that were between 20 Kb and 1 Mb from its nearest promoter were considered to match the distances between interacting cCREs and target genes. Gencode vM4 (GRCm38.p3) was used for protein-coding TSS coordinates. Spearman’s correlation coefficients were calculated for each group and plotted as a function of distance between cCRE and TSS. For genes with alternative TSSs, the distance to the TSS with the highest H3K27ac signal was considered. A spline fit was illustrated using R (smooth.spline function, degree of freedom = 2). Two-sample T-test, two-sided for each 200 Kb interval was performed.

### Clustering analysis of cEnhancer-gene pairs by tissue-/stage-specific H3K27ac

Two groups of cEnhancer-gene pairs were created: 1) pairs with a MAPS-called interaction at E12.5 in one or more tissue type (n=9,370) and 2) pairs with a MAPS-called interaction in one or more stage of developing forebrain (n=11,893). K-means clustering was performed on log2 H3K27ac normalized read counts within 2 Kb of the center of the ATAC peak of cEnhancers. We chose to use 4 main clusters for E12.5 (Fig. 3E) and 5 main clusters for FB (**Fig. S8B**) cEnhancer-gene pairs because adding additional clusters did not reveal new patterns of cEnhancer/gene activity between samples. Sub-clustering was performed for clusters that had high dynamic ranges for signal intensities. Heatmaps were generated using Java TreeView software. Additional sample-specific chromatin features were included. For ChIP-seq data, counts were considered within 2 Kb of ATAC peak center for cEnhancers and 2 Kb of TSS of target genes. Values were quantile normalized and log2 transformed.

### Enhancer validation using VISTA elements and sequence conservation

Validation rates for all cCREs, interacting cCREs, predicted enhancers from Gorkin et al. 2020^7^, and cEnhancers were calculated by determining their ratio of overlapping positive to overlapping negative VISTA elements for in vivo enhancer activity in any tissue. The number of referenced VISTA elements were 1,618 positive and 1,481 negative, which were converted from mm9 to mm10 coordinates using UCSC LiftOver tool (https://genome.ucsc.edu/cgi-bin/hgLiftOver) with default settings. Genes assigned to positive VISTA enhancers were determined for each E12.5 tissue assayed by PLAC-seq. Expression distributions and gene ontology analysis was performed on genes that interacted with a VISTA enhancer element that had positive in vivo activity in the same tissue where a PLAC-seq interaction was observed. Sequence conservation for each element was scored by taking the average PhastCons score for each base pair. PhastConsElements60way and phastConsElements60wayPlacental conserved elements were downloaded from UCSC genome browser. Shuffled control regions were generated using bedtools shuffle with the coordinates of all cCREs as input.

### Gene Ontology and TF Motif enrichment analysis

Gene ontology analysis was performed on gene lists using HOMER findGO.pl function with default parameters. The top enriched GO terms from “Biological process” ontology were reported. TF motif enrichment was performed using HOMER findMotifsGenome.pl function +/- 200bp from cCRE center. Top enriched de novo and known TF motifs were reported for TF gene expression patterns that correlated with TF motif enrichment across tissues.

### GWAS enrichment of cEnhancers and SCZ SNP fine mapping

ATAC peak regions of E12.5 interacting cEnhancers were converted from mm10 to hg19 coordinates using liftover with settings-minMatch 0.5. Regions that did not map back to their original mm10 coordinates were discarded. Using summary statistics for a panel of traits/diseases^73^ we performed linkage disequilibrium score regression on cEnhancers from E12.5 clusters K1, K2, and K3 with all cEnhancers as a local background control set. Schizophrenia (SCZ) 99% credible set SNPs^60^ were leveraged to find cEnhancers overlapping with candidate causal genetic variants.

---

**Figure S1.**
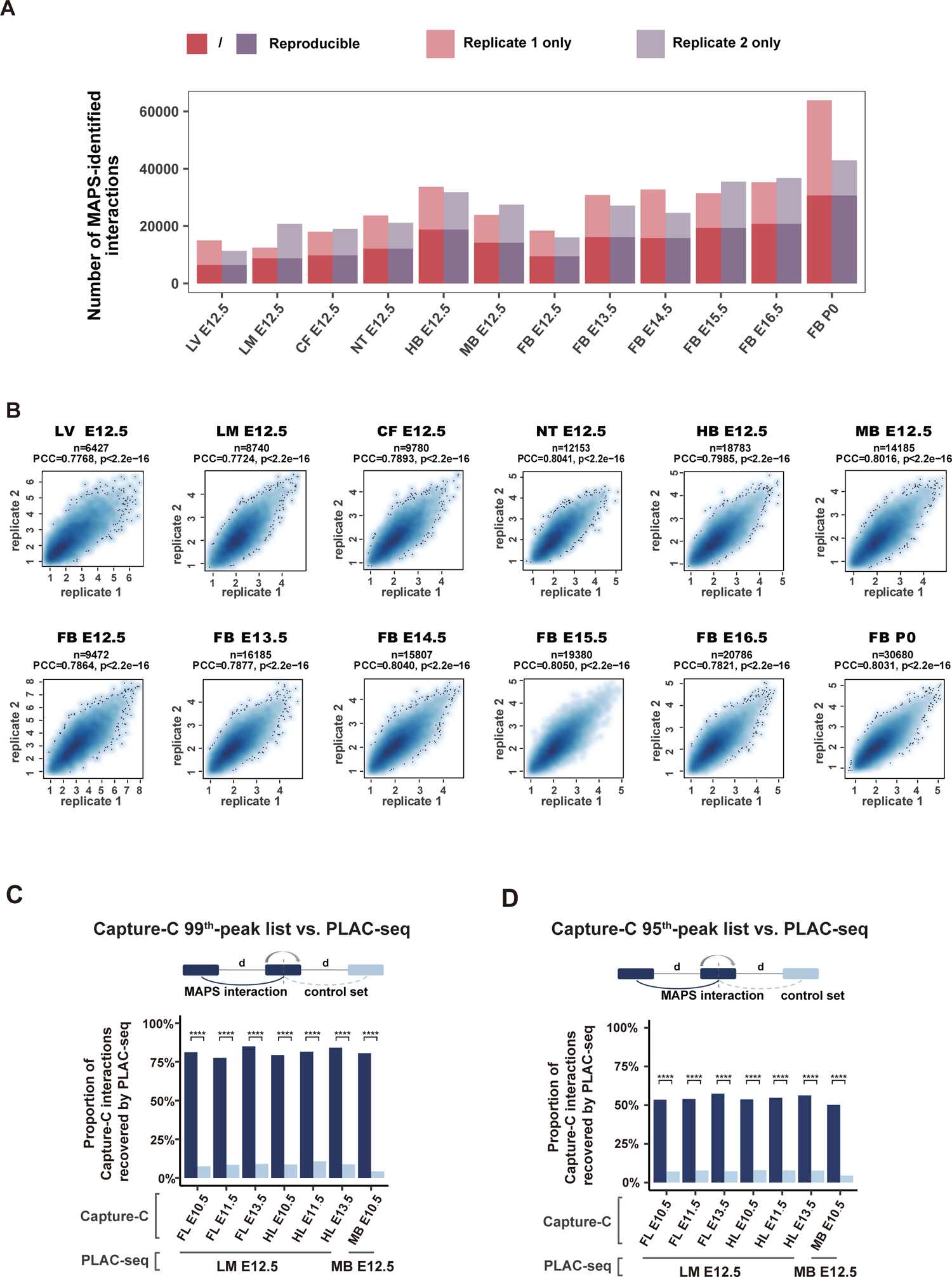
Reproducibility of H3K4me3 PLAC-seq data between two biological replicates. **A.** A bar-plot showing the number of MAPS-identified interactions from each biological replicate with proportions of reproducible interactions highlighted by darker colors. **B.** Scatterplots showing the reproducibility of the normalized contact frequencies between two biological replicates across 12 tissue and stages. Each dot on the plot represents a reproducible interaction and the x- and y-axis correspond to its log2-transformed normalized contact frequencies in two biological replicates. PCC: Pearson correlation coefficients. **C** and **D**. Proportions of interactions identified from published Capture-C datasets^27^ using 99^th^ (**C**) or 95^th^ (**D**) percentile of the empirical distribution as thresholds that overlap with those identified from H3K4me3 PLAC-seq data using the closet tissues (midbrain E10.5 from Capture-C compared with midbrain E12.5 from PLAC-seq; forelimb and hindlimb in Capture-C compared with limb E12.5 from PLAC-seq). Pseudo-interactions that link H3K4me3-containing bin with bin equidistant from but on the other side of H3K4me3-containing bins were used as the control set, as shown in the schematic diagram on the top. The Capture-C interactions with at least one end overlapping reproducible H3K4me3 peaks in limb E12.5 or midbrain E12.5 were considered as testable interactions in this analysis. FL, forelimb. HL, hindlimb. MB, midbrain. E, embryonic day. **** Chi-squared P-value < 2.2e-16.

**Figure S2.**
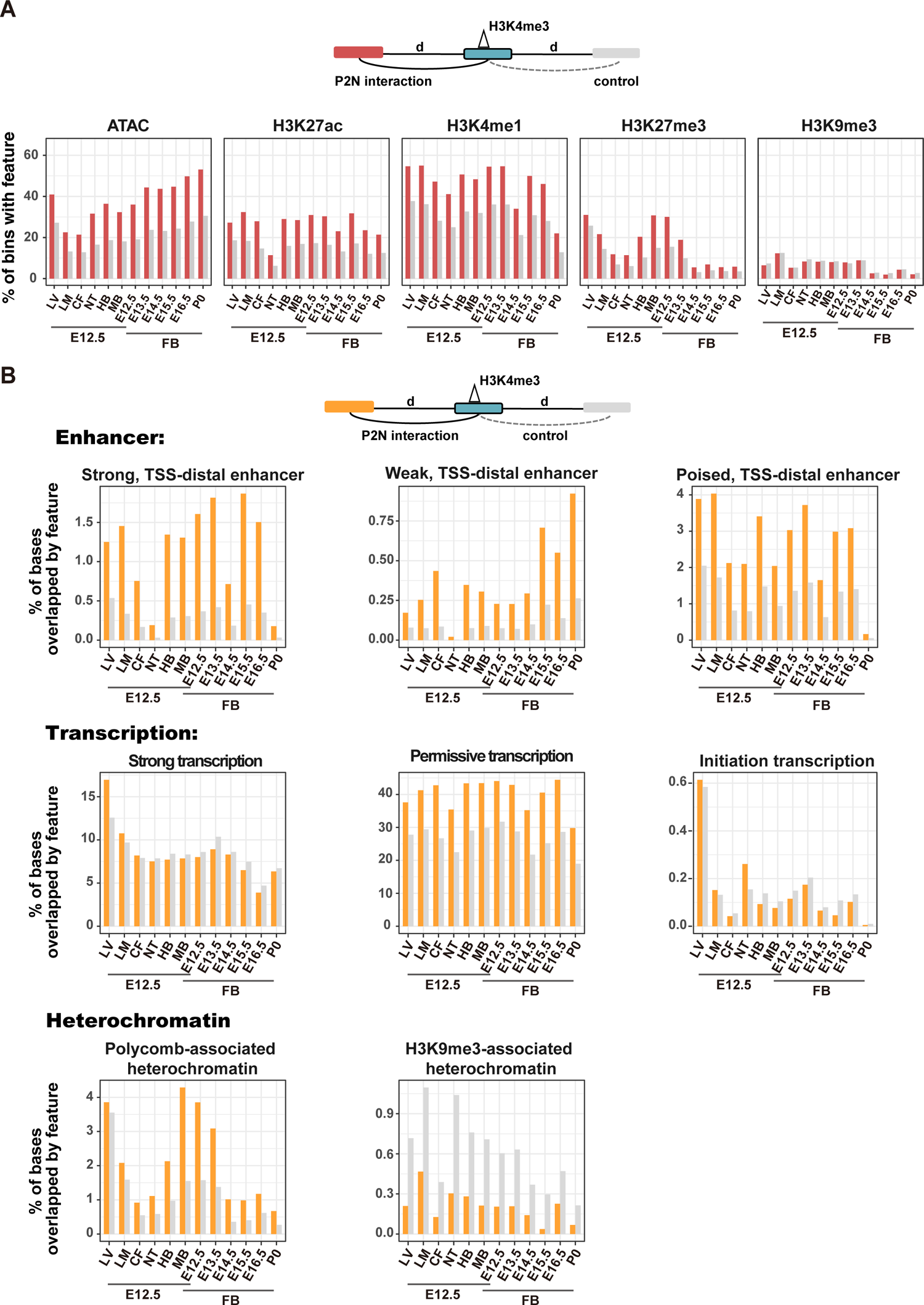
Promoter-interacting regions are significantly enriched for accessible chromatin and specific histone marks across all 12 tissues. **A.** Bar plots showing the percent of promoter-interacting bins overlapping the reproducible peaks identified from ATAC-seq, H3K27ac ChIP-seq, H3K4me1 ChIP-seq, H3K27me3 ChIP-seq or H3K9me3 ChIP-seq from the same tissue. Bins with equidistant from but on the other side of H3K4me3-containing bins were used as the control set, as shown in the schematic diagram on the top. **B.** Bar plots showing the fraction of base pairs in promoter-interacting bins overlapping the reproducible chromHMM state calls from the same tissue. Bins with equidistant from but on the other side of H3K4me3-containing bins were used as the control set, as shown in the schematic diagram on the top.

**Figure S3.**
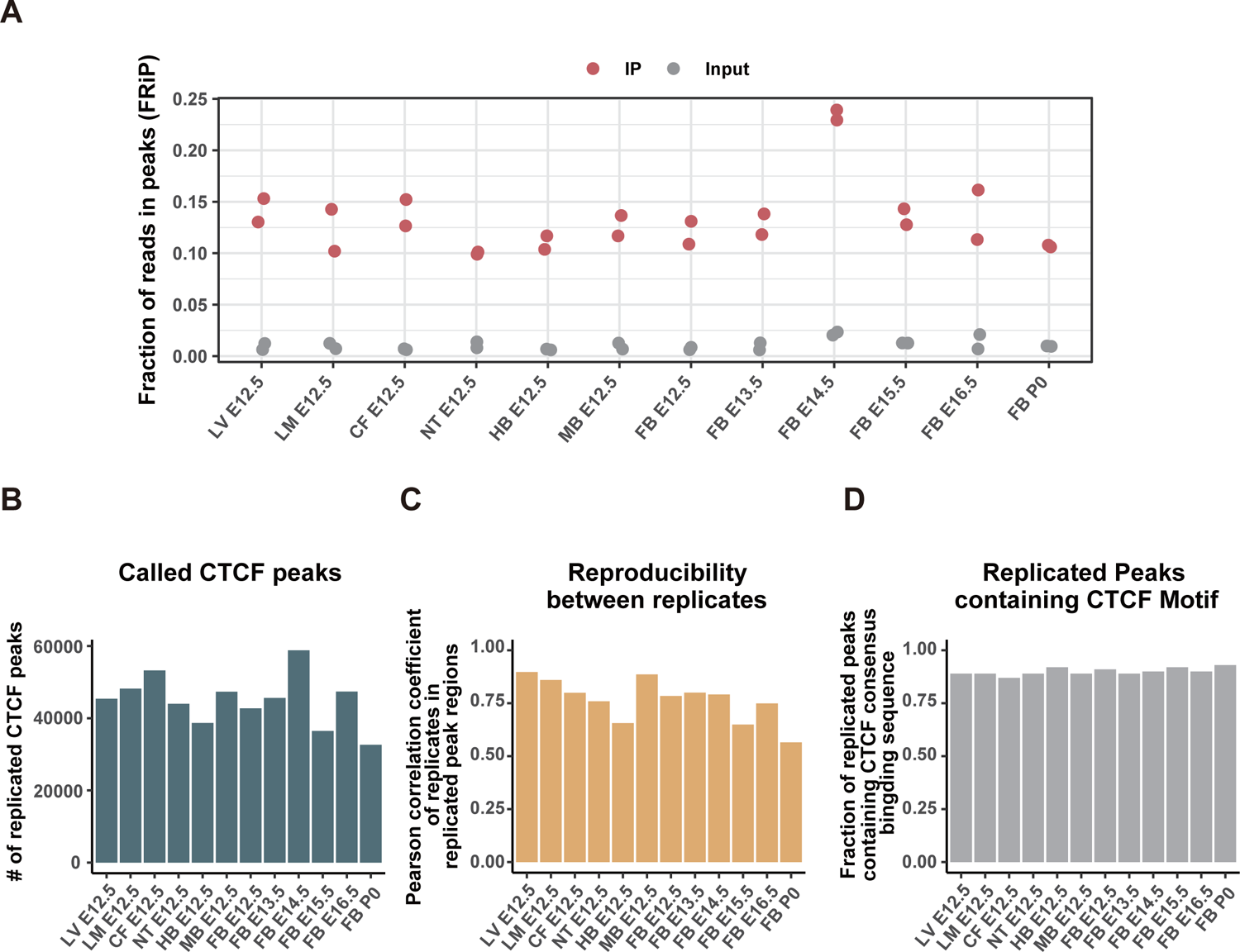
Overview of CTCF ChIP-seq datasets across 12 tissues. **A.** The fraction of all mapped reads in CTCF ChIP-seq libraries and input libraries that fall into the reproducible CTCF ChIP-seq peaks identified by MACS2 (FRiP). Two biological/technical replicates per sample. **B.** Number of replicated peaks identified from CTCF ChIP-seq data across 12 tissues. **C.** Peak reproducibility measured by Pearson correlation of peak strength between biological or technical replicates. **D.** Fraction of replicated peaks containing CTCF consensus binding sequence.

**Figure S4.**
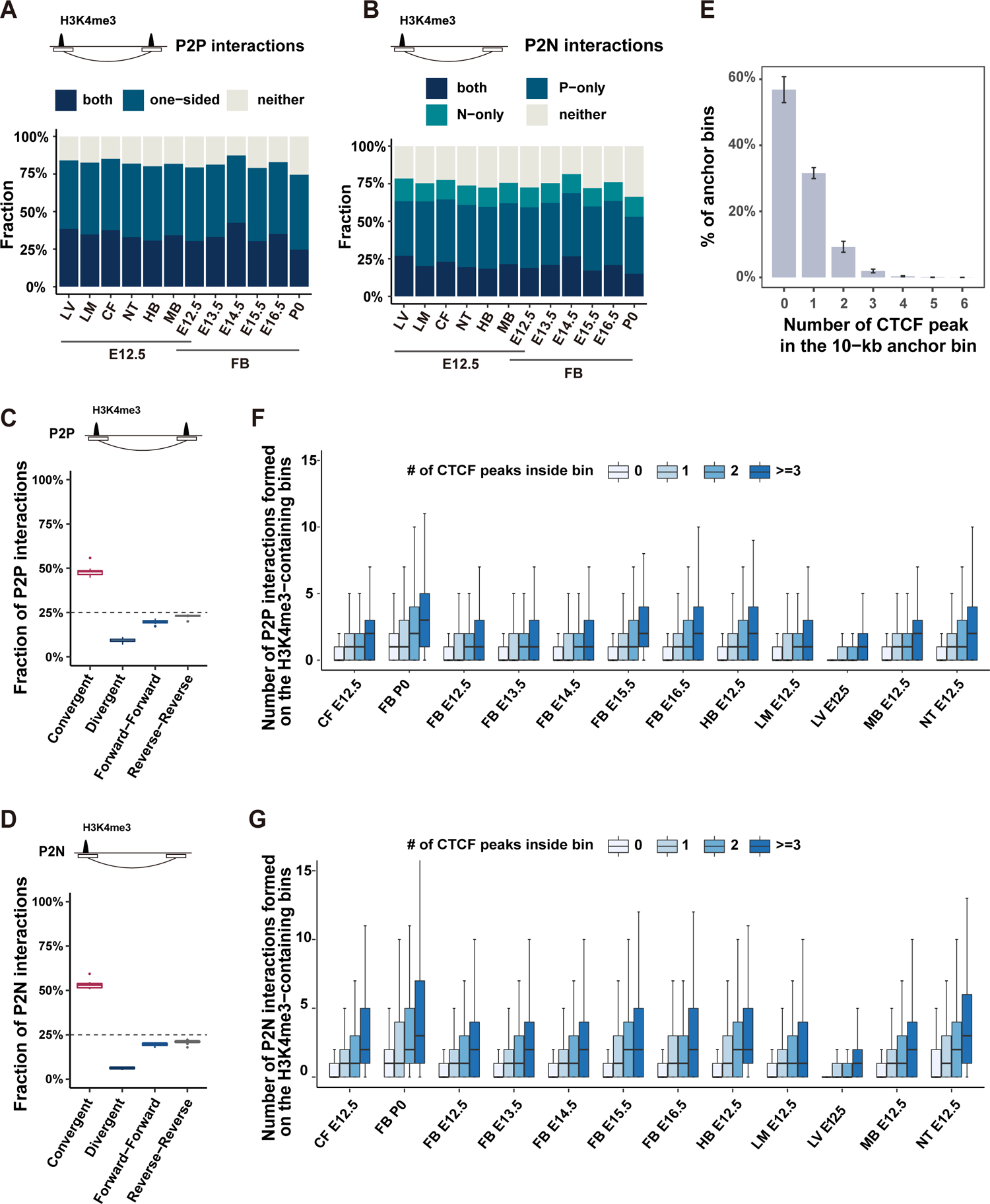
The role of CTCF in promoter-centered chromatin interactions. **A** and **B**. Proportion of long-range P2P (**A**) or P2N (**B**) interactions classified according to presence or absence of CTCF binding on one or both ends in each tissue or developmental stage. P-only: CTCF binding on the “Promoter” end (10 Kb bins with H3K4me3 peak) only; N-only: CTCF binding on the “Non-Promoter” end (10 Kb bins without H3K4me3 peak) only. **C** and **D**. CTCF motif orientation on P2P (**C**) and P2N (**D**) interactions with both ends containing CTCF binding. Only interactions with both ends containing either single CTCF motif or multiple CTCF motifs in the same direction are considered. N=12. **E.** A bar plot showing the distribution of number of CTCF peaks in the H3K4me3-containing bins (anchor bins) in 12 tissue/stage (N=12). The heights of the bars represent the average values of 12 tissues and the error-bars represent the standard deviation. **F** and **G**. Boxplots showing the number of P2P (**F**) or P2N (**G**) interactions formed centered on H3K4me3-containing bins with different numbers of CTCF peaks.

**Figure S5.**
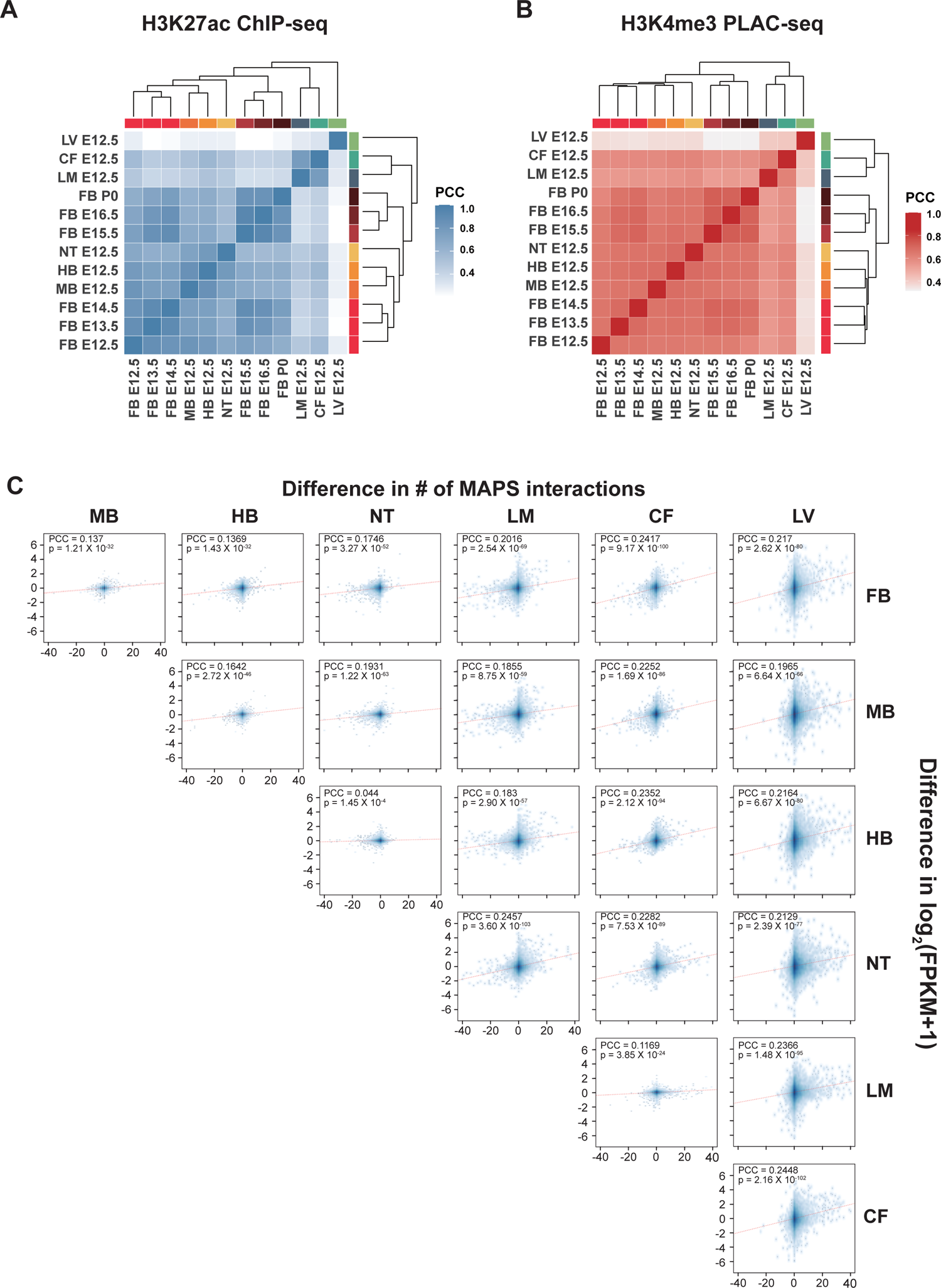
Tissue-to-tissue variability of promoter-centered interactions. **A.** Heatmap showing the Pearson correlation coefficients and hierarchical clustering of H3K27ac ChIP-seq with color bars representing different tissues. **B.** Heatmap showing the Pearson correlation coefficients and hierarchical clustering of normalized contact frequencies identified from combined down-sampled datasets across 12 tissues with color bars representing different tissues. Only interactions detected in at least one tissue are considered in this analysis. **C.** Similar to Fig 2C, scatterplot between the difference of the number of MAPS-identified significant chromatin interactions (x-axis) and the different of gene expression (y-axis, merged by log2(FPKM+1)), for all 21 pairs of 7 tissues at e12.5. The red dashed line represents the fitted linear line, suggesting that the change of significant chromatin interactions is positively correlated with the change of gene expression.

**Figure S6.**
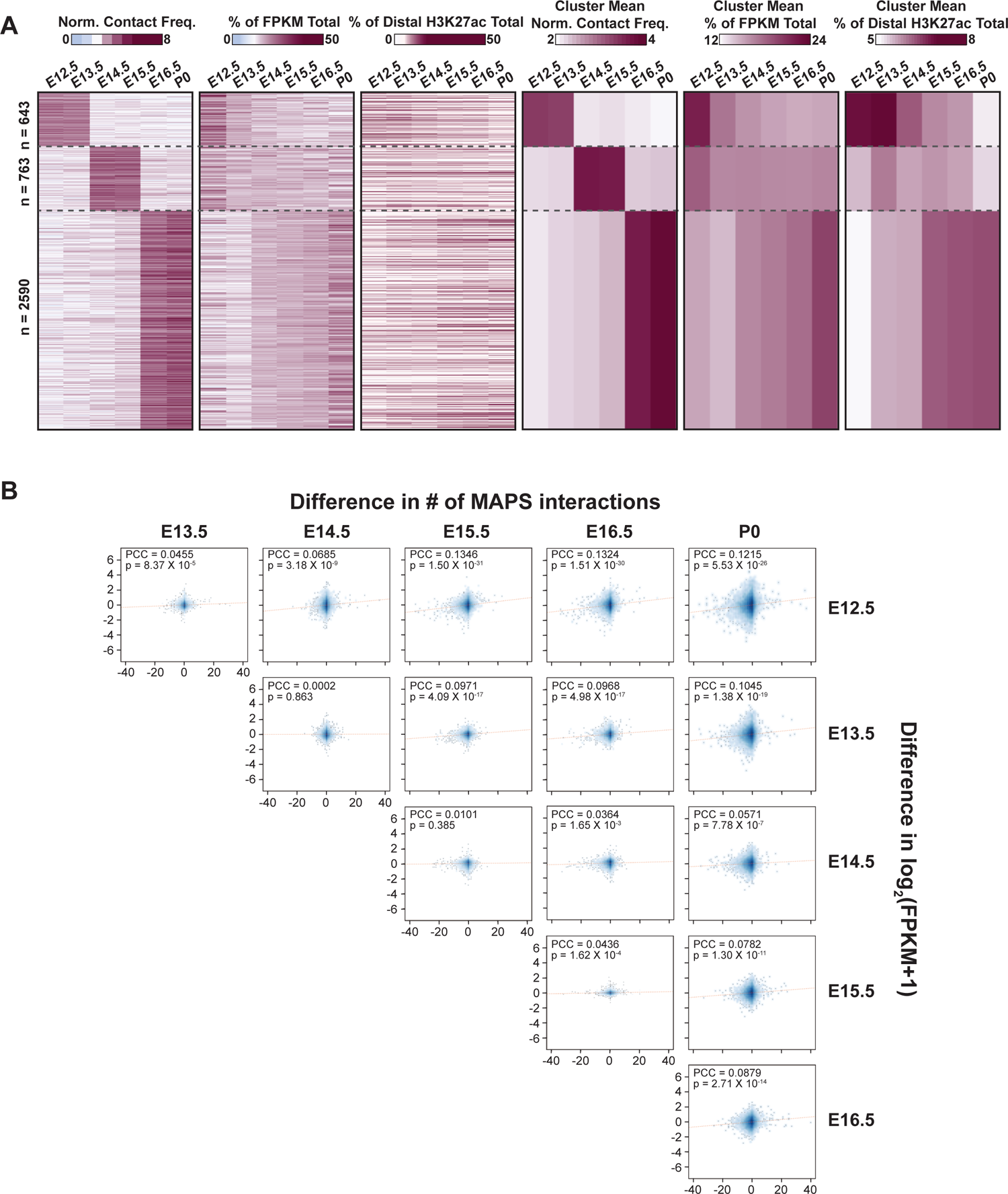
Dynamics of promoter-centered interactions across developmental stages in forebrain. **A.** Heatmap displaying normalized contact frequencies, gene expression of interacting promoters, and H3K27ac distal peak signal in peak to non-peak stage-specific interactions. Gene expression and H3K27ac are represented as individual tissue percentage of total sum across all ti ssues. Tissue-specific interactions are clustered for being exclusive in E12.5/E13.5, E14.5/E15.5, or E16.5/P0-only interactions. **B.** Similar to Fig. 2C, scatterplot between the difference of the number of MAPS-identified significant chromatin interactions (x-axis) and the different of gene expression (y-axis, merged by log2(FPKM+1)), for all 15 pairs of forebrain tissue at 6 developmental stages. The red dashed line represents the fitted linear line, suggesting that the change of significant chromatin interactions is positively correlated with the change of gene expression.

**Figure S7.**
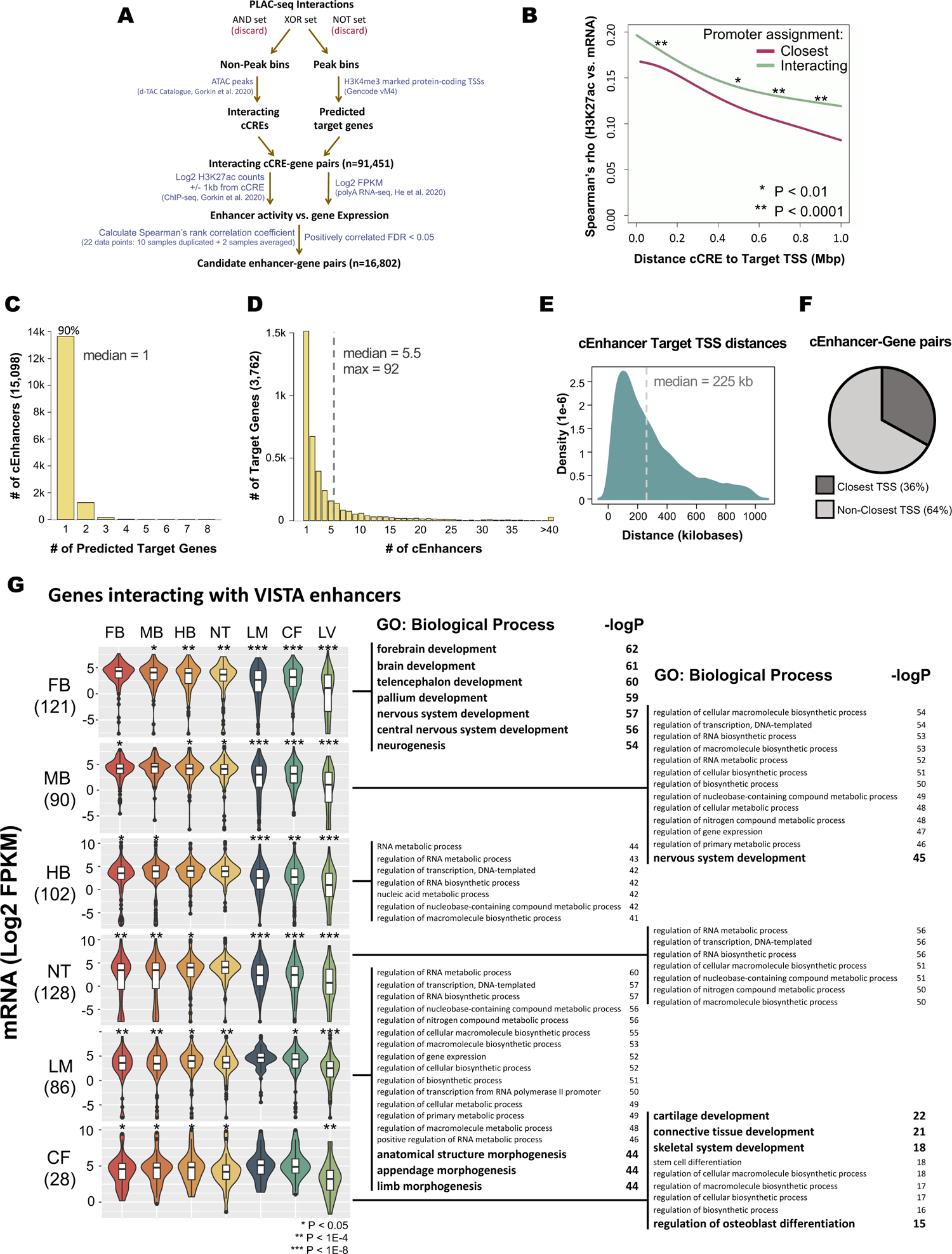
Chromatin interactions identify enhancer target genes. **A.** Schematic illustrating analysis pipeline for determining interacting cCRE-gene pairs and cEnhancer-gene pairs. **B.** Spline fit lines of Spearman’s correlation coefficients between cCRE H3K27ac and assigned gene mRNA as a function of cCRE to TSS distance for two different pairing strategies. Degrees of freedom = 2, Two-sample T-test, two-sided for each 200 Kb interval. **C.** Histogram for number of target genes assigned to cEnhancers. **D.** Histogram for number of cEnhancers assigned to target genes. **E.** Density plot of cEnhancer to target TSS distances. **F.** Pie chart representing percentage of pairs where the cEnhancer targets the gene with the closest TSS or not. **G.** Expression and top enriched GO: Biological Process terms for genes interacting with VISTA enhancers with an identified promoter interaction in the same tissue where enhancer activity was reported positive in the VISTA database. Due to low numbers of liver positive VISTA enhancers, only one gene (*Igf2*) was assigned to this group and was omitted from the figure. Paired T-test, two-sided.

**Figure S8.**
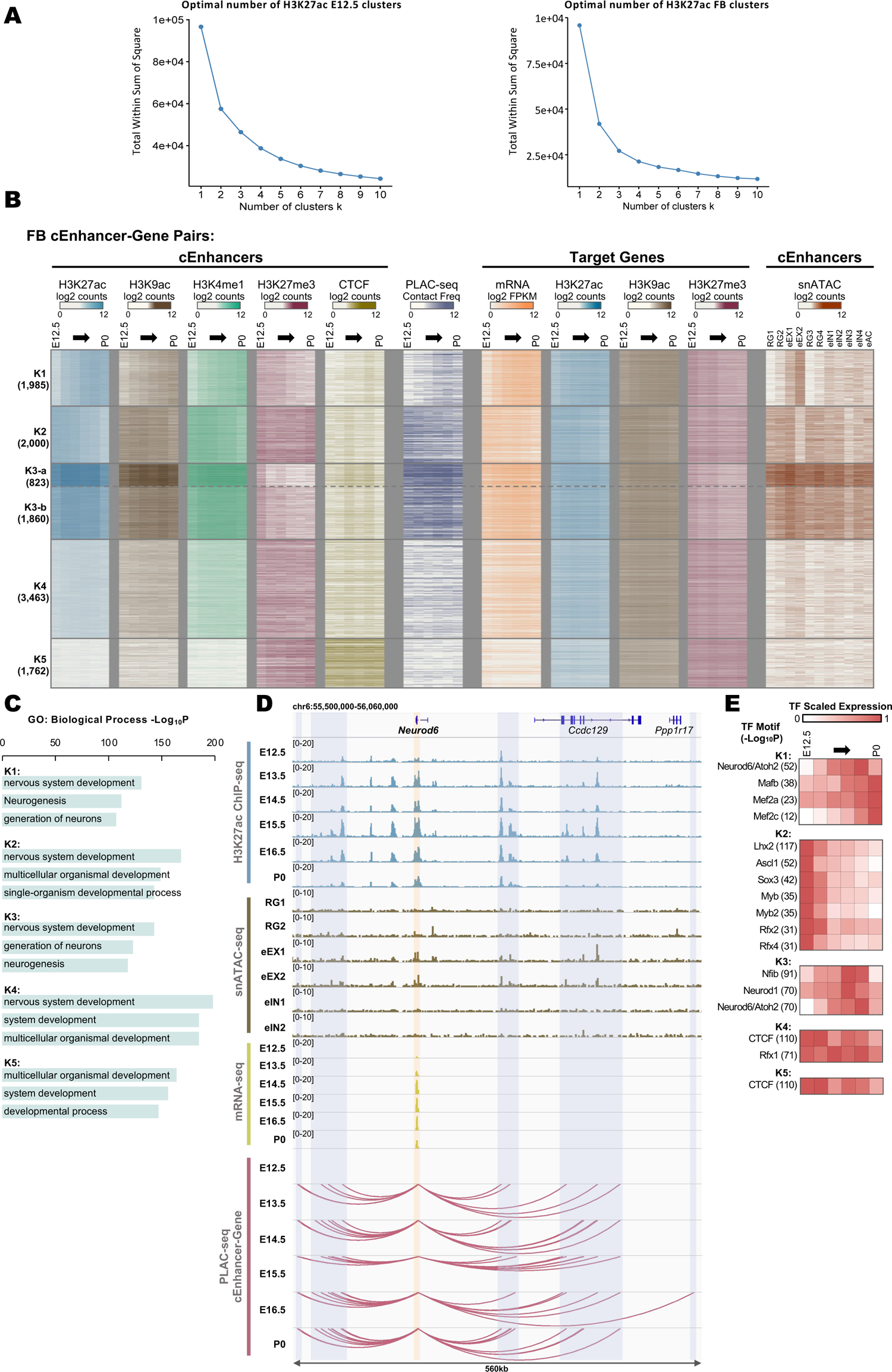
Profiling candidate forebrain enhancer-gene pairs across developmental stages. **A.** Elbow plots displaying sum of squared distances for indicated number of kmeans clusters. Kmeans clustering was performed using normalized H3K27ac counts surrounding 2 Kb of cEnhancers of cEnhancer-gene pairs present in E12.5 tissues (left) and FB developmental stages (right). **B.** Heatmap for chromatin features and expression of forebrain cEnhancer-gene pairs. Pairs were k-means clustered by H3K27ac signal surrounding 2 Kb at center of cEnhancer. **C.** Top enriched GO: Biological Process terms for genes of clusters from **Fig. S8B**. **D.** Example gene, *Neurod6*, showing correlated stage-specific cEnhancer H3K27ac, gene expression, and cEnhancer-gene interactions. The cEnhancer-gene interactions in this region from FB at different stages were marked by arcs. *Neurod6* TSS region was highlighted by yellow box and the cEnhancers are highlighted by blue boxes. **E.** Enriched known and *de novo* motifs that have TF gene expression matching cEnhancer H3K27ac patterns across stages.

**Figure S9.**
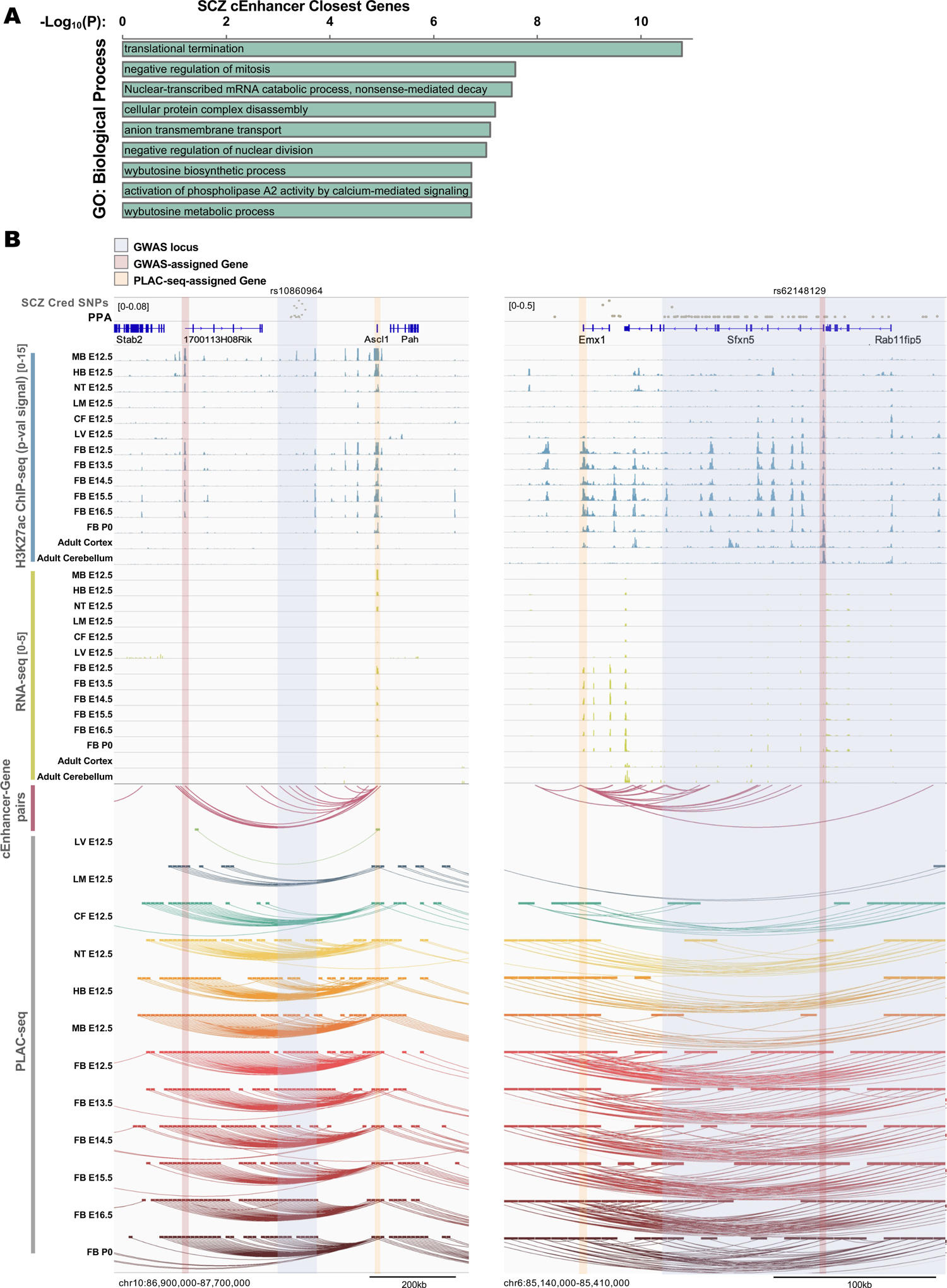
SCZ-associated non-coding variants are enriched at fetal-specific enhancers. **A.** Gene ontology analysis showing top enriched Biological Process terms of genes closest to cEnhancers harboring SCZ credible set SNPs by TSS proximity at 1D genomic distance. **B.** SCZ example genes, *Ascl1* and *Emx1*, showing correlated tissue-specific cEnhancer H3K27ac, gene expression, and chromatin interactions at loci harboring SCZ SNPs.

**Supplementary Table 1.** Information about each PLAC-seq library.

| Sample name | # total reads | # usable reads | % inter-chromosomal (*trans*) reads | % intra-chromosomal (*cis*) reads over 1Kb | Fraction of reads in peaks (FRiP) |
| --- | --- | --- | --- | --- | --- |
| CF E12.5 replicate1 | 205,620,027 | 114,214,116 | 35.4% | 78.6% | 9.5% |
| CF E12.5 replicate2 | 205,491,553 | 120,760,796 | 35.9% | 79.2% | 9.8% |
| HB E12.5 replicate1 | 199,944,273 | 119,464,746 | 33.7% | 78.4% | 13.4% |
| HB E12.5 replicate 2 | 292,213,527 | 181,478,442 | 31.3% | 75.4% | 7.9% |
| LM E12.5 replicate1 | 248,569,504 | 136,610,243 | 28.6% | 84.8% | 12.3% |
| LM E12.5 replicate2 | 217,661,328 | 120,652,338 | 26.2% | 82.9% | 16.6% |
| LV E12.5 replicate1 | 290,025,435 | 173,471,834 | 30.3% | 80.2% | 10.8% |
| LV E12.5 replicate2 | 209,971,157 | 129,280,407 | 27.0% | 75.6% | 14.4% |
| MB E12.5 replicate1 | 194,282,599 | 118,233,276 | 31.6% | 79.8% | 10.6% |
| MB E12.5 replicate2 | 274,285,709 | 160,942,810 | 34.9% | 80.5% | 8.7% |
| NT E12.5 replicate1 | 234,384,550 | 124,201,120 | 21.4% | 55.8% | 11.9% |
| NT E12.5 replicate2 | 160,248,906 | 93,456,174 | 27.7% | 83.9% | 10.4% |
| FB E12.5 replicate 1 | 202,525,962 | 130,620,129 | 28.3% | 74.6% | 10.4% |
| FB E12.5 replicate 2 | 206,487,951 | 138,052,881 | 23.2% | 52.8% | 12.1% |
| FB E13.5 replicate 1 | 330,937,332 | 178,556,964 | 35.1% | 72.9% | 10.7% |
| FB E13.5 replicate 2 | 281,352,866 | 146,882,201 | 29.3% | 67.6% | 11.5% |
| FB E14.5 replicate 1 | 351,516,759 | 199,453,133 | 30.2% | 71.6% | 7.8% |
| FB E14.5 replicate 2 | 183,756,702 | 111,749,314 | 27.7% | 75.0% | 13.5% |
| FB E15.5 replicate 1 | 161,356,716 | 101,989,803 | 32.6% | 80.0% | 16.2% |
| FB E15.5 replicate 2 | 195,232,542 | 123,484,382 | 28.1% | 78.5% | 11.3% |
| FB E16.5 replicate 1 | 198,167,464 | 137,340,506 | 27.7% | 74.3% | 13.9% |
| FB E16.5 replicate 2 | 169,762,737 | 102,678,742 | 24.8% | 76.9% | 15.0% |
| FB P0 replicate 1 | 207,017,793 | 119,576,342 | 25.9% | 84.8% | 15.1% |
| FB P0 replicate 2 | 171,494,682 | 100,886,728 | 24.6% | 80.2% | 13.8% |

**Supplementary Table 2.**
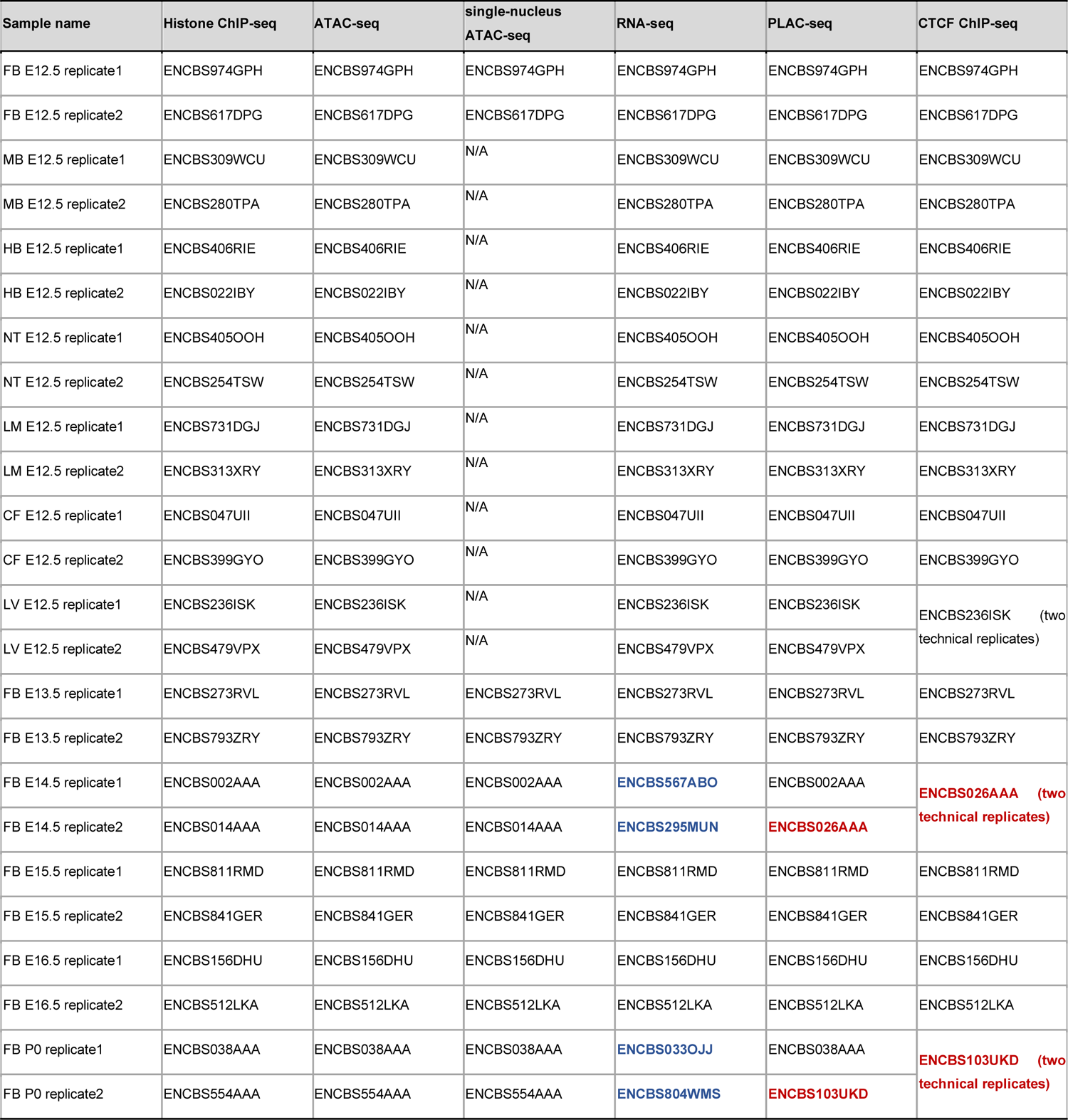
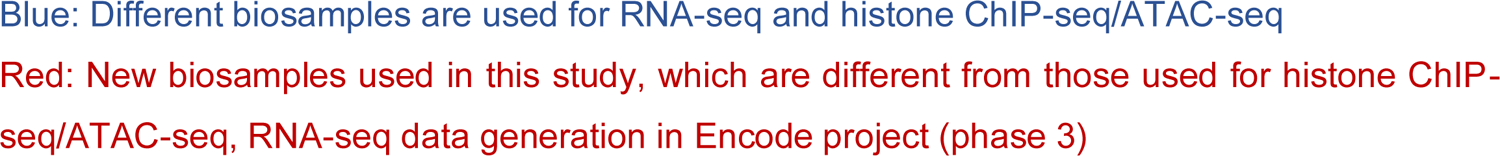
Information about the biological material used as input material for each experimental assay. The biosample accession information of the histone ChIP-seq, RNA-seq and ATAC-seq datasets was extracted from ENCODE portal (https://www.encodeproject.org/). For the PLAC-seq and CTCF ChIP-seq datasets generated in this study, the biosample information was provided, with majority of them using the same biosamples as histone ChIP-seq, RNA-seq or ATAC-seq. The exceptions are marked by dark red.

| Sample name | Histone ChIP-seq | ATAC-seq | single-nucleus ATAC-seq | RNA-seq | PLAC-seq | CTCF ChIP-seq |
| --- | --- | --- | --- | --- | --- | --- |
| FB E12.5 replicate1 | ENCBS974GPH | ENCBS974GPH | ENCBS974GPH | ENCBS974GPH | ENCBS974GPH | ENCBS974GPH |
| FB E12.5 replicate2 | ENCBS617DPG | ENCBS617DPG | ENCBS617DPG | ENCBS617DPG | ENCBS617DPG | ENCBS617DPG |
| MB E12.5 replicate1 | ENCBS309WCU | ENCBS309WCU | N/A | ENCBS309WCU | ENCBS309WCU | ENCBS309WCU |
| MB E12.5 replicate2 | ENCBS280TPA | ENCBS280TPA | N/A | ENCBS280TPA | ENCBS280TPA | ENCBS280TPA |
| HB E12.5 replicate1 | ENCBS406RIE | ENCBS406RIE | N/A | ENCBS406RIE | ENCBS406RIE | ENCBS406RIE |
| HB E12.5 replicate2 | ENCBS022IBY | ENCBS022IBY | N/A | ENCBS022IBY | ENCBS022IBY | ENCBS022IBY |
| NT E12.5 replicate1 | ENCBS405OOH | ENCBS405OOH | N/A | ENCBS405OOH | ENCBS405OOH | ENCBS405OOH |
| NT E12.5 replicate2 | ENCBS254TSW | ENCBS254TSW | N/A | ENCBS254TSW | ENCBS254TSW | ENCBS254TSW |
| LM E12.5 replicate1 | ENCBS731DGJ | ENCBS731DGJ | N/A | ENCBS731DGJ | ENCBS731DGJ | ENCBS731DGJ |
| LM E12.5 replicate2 | ENCBS313XRY | ENCBS313XRY | N/A | ENCBS313XRY | ENCBS313XRY | ENCBS313XRY |
| CF E12.5 replicate1 | ENCBS047UII | ENCBS047UII | N/A | ENCBS047UII | ENCBS047UII | ENCBS047UII |
| CF E12.5 replicate2 | ENCBS399GYO | ENCBS399GYO | N/A | ENCBS399GYO | ENCBS399GYO | ENCBS399GYO |
| LV E12.5 replicate1 | ENCBS236ISK | ENCBS236ISK | N/A | ENCBS236ISK | ENCBS236ISK | ENCBS236ISK (two technical replicates) |
| LV E12.5 replicate2 | ENCBS479VPX | ENCBS479VPX | N/A | ENCBS479VPX | ENCBS479VPX |  |
| FB E13.5 replicate1 | ENCBS273RVL | ENCBS273RVL | ENCBS273RVL | ENCBS273RVL | ENCBS273RVL | ENCBS273RVL |
| FB E13.5 replicate2 | ENCBS793ZRY | ENCBS793ZRY | ENCBS793ZRY | ENCBS793ZRY | ENCBS793ZRY | ENCBS793ZRY |
| FB E14.5 replicate1 | ENCBS002AAA | ENCBS002AAA | ENCBS002AAA | ENCBS567ABO | ENCBS002AAA | ENCBS026AAA (two technical replicates) |
| FB E14.5 replicate2 | ENCBS014AAA | ENCBS014AAA | ENCBS014AAA | ENCBS295MUN | ENCBS026AAA |  |
| FB E15.5 replicate1 | ENCBS811RMD | ENCBS811RMD | ENCBS811RMD | ENCBS811RMD | ENCBS811RMD | ENCBS811RMD |
| FB E15.5 replicate2 | ENCBS841GER | ENCBS841GER | ENCBS841GER | ENCBS841GER | ENCBS841GER | ENCBS841GER |
| FB E16.5 replicate1 | ENCBS156DHU | ENCBS156DHU | ENCBS156DHU | ENCBS156DHU | ENCBS156DHU | ENCBS156DHU |
| FB E16.5 replicate2 | ENCBS512LKA | ENCBS512LKA | ENCBS512LKA | ENCBS512LKA | ENCBS512LKA | ENCBS512LKA |
| FB P0 replicate1 | ENCBS038AAA | ENCBS038AAA | ENCBS038AAA | ENCBS033OJJ | ENCBS038AAA | ENCBS103UKD (two technical replicates) |
| FB P0 replicate2 | ENCBS554AAA | ENCBS554AAA | ENCBS554AAA | ENCBS804WMS | ENCBS103UKD |  |

Blue: Different biosamples are used for RNA-seq and histone ChIP-seq/ATAC-seq
Red: New biosamples used in this study, which are different from those used for histone ChIP-seq/ATAC-seq, RNA-seq data generation in Encode project (phase 3)

**Supplementary Table 3.** The identifiers of processed files downloaded from ENCODE data portal (https://www.encodeproject.org/).

